# Erotic cue exposure increases physiological arousal, biases choices towards immediate rewards and attenuates model-based reinforcement learning

**DOI:** 10.1101/2022.09.04.506507

**Authors:** David Mathar, Annika Wiebe, Deniz Tuzsus, Kilian Knauth, Jan Peters

## Abstract

Computational psychiatry focuses on identifying core cognitive processes that appear altered across a broad range of psychiatric disorders. Temporal discounting of future rewards and model-based control during reinforcement learning have proven as two promising candidates. Despite its trait-like stability, temporal discounting has been suggested to be at least partly under contextual control. For example, highly arousing cues such as erotic pictures were shown to increase discounting, although overall evidence to date remains somewhat mixed. Whether model-based reinforcement learning is similarly affected by arousing cues is unclear. Here we tested cue-reactivity effects (erotic pictures) on subsequent temporal discounting and model-based reinforcement learning in a within-subjects design in n=39 healthy male participants. Self-reported and physiological arousal (cardiac activity and pupil dilation) were assessed before and during cue exposure. Arousal was increased during exposure of erotic vs. neutral cues both on the subjective and autonomic level. Erotic cue exposure nominally increased discounting as reflected by reduced choices of delayed options. Hierarchical drift diffusion modeling (DDM) linked increased discounting to a shift in the starting point bias of evidence accumulation towards immediate options. Model-based control during reinforcement learning was reduced following erotic cues according to model-agnostic analysis. Notably, DDM linked this effect to attenuated forgetting rates of unchosen options, leaving the model-based control parameter unchanged. Our findings replicate previous work on cue-reactivity effects in temporal discounting and for the first time show similar effects in model-based reinforcement learning. Our results highlight how environmental cues can impact core human decision processes and reveal that comprehensive drift diffusion modeling approaches can yield novel insights in reward-based decision processes.

## Introduction

Decisions often entail consequences that differ in temporal proximity and value. The tendency to favor smaller-but-sooner over larger-but-later rewards, known as temporal discounting, is common across animals (Kalenscher & Pennartz, 2008) and humans (Peters & Büchel, 2011). The individual degree of temporal discounting varies profoundly and appears stable across time (Simpson & Vuchinich, 2000; Arfer & Luhmann, 2017; Kirby, 2009) and across different testing environments (Bruder et al., 2021; Odum, 2011). Thus, discounting is regarded as a trait-like characteristic (Smith & Hantula, 2008). Notably, alterations in temporal discounting are associated with a range of psychiatric disorders, including substance use disorders, gambling disorder, obesity and attention-deficit hyperactivity disorder (Amlung et al., 2016, 2019; Bickel et al., 2019; Jackson & MacKillop, 2016; Wiehler & Peters, 2015).

Despite its trait-like stability, temporal discounting is at least partly under contextual control (Lempert et al., 2016; Peters & Büchel, 2011). Temporal discounting is attenuated when future reward options are paired with participant-specific episodic cues (Bromberg et al., 2017; Peters & Büchel, 2010; Rösch et al., 2022). In contrast, exposure to highly arousing cues such as erotic pictures can increase temporal discounting (Kim & Zauberman, 2013; Van den Bergh et al., 2008; Wilson & Daly, 2004). However, findings regarding effects of arousing cues on temporal discounting appear somewhat heterogenous, with two more recent studies reporting no significant effect of erotic cue exposure on the degree of temporal discounting (Simmank et al., 2015; Knauth & Peters, 2022). Notably, both studies implemented cue exposure in an event-related manner during temporal discounting in contrast to earlier studies that used blocked designs in which participants were first exposed to multiple arousing pictures and subsequently completed the discounting task (Kim & Zauberman, 2013; Van den Bergh et al., 2008; Wilson & Daly, 2004).

Besides differing in their short-term and long-term consequences, decisions can arise from both habits and goal-directed planning. This competition for behavioral control has been operationalized in reinforcement learning models in terms of model-free and model-based learning strategies. Learning from previous experience is essential for optimal behavior in volatile environments and is tightly linked to reward prediction errors signaled by dopaminergic midbrain neurons (Schultz, Dayan & Montague, 1997). This reliance on stimulus-reward associations is referred to as model-free reinforcement learning. In contrast, model-based reinforcement learning is related to a computationally more demanding incorporation of a cognitive model of the environment during learning to facilitate goal-directed action selection (Balleine & O’Doherty, 2010; Daw et al., 2011; Dolan & Dayan, 2013). The relative degree of model-based as compared to model-free reinforcement learning has been linked to striatal dopamine transmission (Deserno, 2015). More importantly, model-based control might constitute a trans-diagnostic marker across distinct psychopathologies (Culbreth et al., 2016; Gillan et al., 2016; Wyckmans et al., 2019), similar to corresponding observations in temporal discounting (Amlung et al., 2016; Jackson & MacKillop, 2016; Bickel et al., 2019). First evidence that model-based reinforcement learning can be modulated stems from pharmacological interventions that boost catecholamine transmission. L-Dopa intake was reported to increase model-based control (Sharp et al., 2016; Wunderlich et al., 2012). However, this finding was challenged recently (Deserno et al., 2021; Kroemer et al., 2019). In recent own work, we found that a single dose of the catecholamine precursor L-Tyrosine enhances model-based reinforcement learning to some extend in healthy participants (Mathar et al., 2022). However, whether model-based control is also partly under contextual control in a similar manner to temporal discounting, is still elusive. Patzelt et al., (2019) reported that increasing incentive magnitude can increase model-based control in a large online cohort. We also found first evidence for modulatory effects of model-based control by context. Participants suffering from gambling disorder showed enhanced model-based reinforcement learning in a highly arousing gambling environment compared with task performance in a neutral café (Wagner et al., 2022). Yet, no work exists that examined potential modulatory effects of arousing cues such as erotic pictures on model-based reinforcement learning in healthy participants.

Why would erotic cues affect learning and decision-making? Erotic cues robustly activate neural structures including ventral striatum and orbitofrontal cortex that receive catecholaminergic projections from midbrain nuclei (Gola et al., 2016; Stark et al., 2005; Wehrum-Osinsky et al., 2014). Via this route, primary reinforcers such as erotic cues are assumed to promote immediate out-of-domain reward preferences (Li, 2008; Yeomans & Grace, 2015) and thus might also promote habitual (model-free) over goal-directed (model-based) control of behavior. Effects of such cues on decision-making in healthy individuals may also share conceptual similarities with so-called cue-reactivity responses in individuals suffering from addiction. Cue-reactivity effects in substance use-disorders and behavioral addictions reflect increased arousal on the subjective, physiological, and neural level in response to addiction-related cues (Courtney et al., 2015; Starcke et al., 2018; Volkow et al., 2010). Ascending catecholaminergic projections originating in the brainstem arousal systems may partly drive these cue effects, as salient stimuli are associated with firing of neurons in the locus coeruleus, the primary noradrenergic brainstem nucleus (Bouret & Richmond, 2015; Chen & Sara, 2007; Mather et al., 2016). This facilitates noradrenaline release in widely distributed subcortical and cortical regions resulting in enhanced stimulus processing (Howells et al., 2012; Mather et al., 2016; Vazey et al., 2018). Locus coeruleus activity is tightly linked to pupil dilation (Aston-Jones & Cohen, 2005; Joshi et al., 2016) in primates (Varazzani et al., 2015) as well as humans (Murphy et al., 2014), although this relationship is strongly influenced by brain states (Megemont et al. 2022). In line with this, pupil dilation is increased following erotic cue exposure (Rieger et al., 2012; Finke et al., 2017). Besides affecting pupil dilation, arousal related noradrenaline release also modulates heart rate and heart rate variability (Bradley et al., 2008; Schneider et al., 2016; Gordan et al., 2015).

Here we examined the effect of block-wise erotic versus neutral cue exposure on temporal discounting and model-based reinforcement learning in a within-subject approach. Importantly, we expanded upon previous work in several ways. First, there is no study to date that assessed erotic cue exposure on model-based reinforcement learning. Second, we utilized a combination of temporal discounting and reinforcement learning models with a drift diffusion model-based choice rule (Fontanesi et al., 2019; Pedersen et al., 2017; Peters & D’Esposito, 2020; Shahar et al., 2019; Wagner et al., 2020, 2022). This enables a more comprehensive account of participants’ learning and decision-making behavior by decomposing observed RT distributions and corresponding choices into latent underlying processes. Third, we comprehensively assessed effects of erotic cue exposure on both self-reported and physiological markers of arousal, specifically spontaneous eye blink rate, pupil dilation, heart rate and heart rate variability, thereby advancing over previous studies that focused solely on behavioral measures (Kim & Zauberman, 2013; Van den Bergh et al., 2008; Wilson & Daly, 2004). Arousal measures were obtained both before and during cue exposure. We assumed that highly arousing (erotic) cues may modulate temporal discounting and model-based control via promotion of short-term reward preferences in the light of enhanced autonomic arousal.

## Methods

### Participants

Forty healthy male volunteers (right-handed, non-smoking, no history of psychiatric or neurological illness, no medication or drug use), participated in the study. The sample size was calculated in an a-priori power analysis based on findings of erotic cue exposure on temporal discounting by Wilson & Daly (2004) via G*Power 3.1 (Faul, Erdfelder, Buchner, & Lang, 2007). We estimated a target sample size of n=39 with an estimated effect size d=0.6, α=0.05 and power β=0.95. All volunteers provided written informed consent and received a reimbursement of 10 € per hour for their participation. Performance-dependent additional reimbursement is outlined in the task specific sections further below. The study was approved by the ethics committee of the University of Cologne Medical Center. One participant was excluded prior to data analysis due to inattentiveness during cue exposure.

### General Procedure

Participants visited our lab twice on two separate days (mean distance [range]: 6 [1-22] days). On each testing day, participants first underwent a baseline physiological assessment of five minutes (see below). Subsequently, they performed a first cue exposure block. In each cue exposure block, a series of 21 erotic or neutral pictures was presented on a computer screen twice in two separate rounds and participants rated each picture first according to arousal and second according to valence. Then participants performed a temporal discounting task. After task completion participants executed a second cue exposure block of 21 (10 new, 11 already rated) neutral or erotic pictures and rated them again according to arousal and valence similar to the first cue exposure block. Finally, they performed a sequential reinforcement learning task to assess the individual degree of model-free and model-based control over behavior.

Below, physiological assessment, cue exposure and tasks are delineated in more detail. Physiological data assessment and task implementations are similar to those reported in Mathar et al. (2022).

### Physiological data acquisition

For quantification of possible effects of erotic cue exposure on physiological arousal, we assessed three different measures of autonomic nervous system activity, spontaneous eye blink rate, heart rate, and pupil dilation. Measurements were taken for five minutes prior to cue exposure and during both cue exposure blocks (erotic and neutral) before temporal discounting and sequential reinforcement learning tasks were performed.

Spontaneous eye blink rate is discussed as a proxy measure for central catecholamine levels (Elsworth et al., 1991; Groman et al., 2014; Kaminer et al., 2011; Sescousse et al., 2018). Heart rate and pupil dilation are tightly linked to the interplay of sympathetic and parasympathetic afferents (Berntson et al., 1997; Bradley et al., 2008; Schneider et al., 2016) and arousal-related noradrenaline transmission (Gordan et al., 2015; Murphy et al., 2014; Phillips et al., 2000; Preuschoff et al., 2011). In addition, we computed the low frequency-to-high frequency (LF/HF) heart rate variability ratio as an index of the balance between sympathetic and parasympathetic activity. A relative increase in the HF variability component indicates enhanced parasympathetic input, whereas increased variability in the LF component indicates stronger sympathetic input (Appelhans & Luecken, 2008; Pumprla et al., 2002). There is evidence, that LF/HF ratio is modulated by salient cues and might covary with substance related craving and relapse risk (Garland et al., 2012; Witteman et al., 2015).

In each of the three physiological assessments per testing day, participants were seated in a shielded, dimly lit room 0.6m from a 24-inch LED screen (resolution: 1366 × 768 px; refresh rate: 60 Hz) with their chin and forehead placed in a height-adjustable chinrest. Stimulus presentation was implemented using the Psychophysics toolbox (Version 3.0.14) for MATLAB (R2017a; MathWorks, Natick, MA). Participants were instructed to move as little as possible and fixate a white cross on a grey background presented on the screen. Spontaneous eye blink rate was recorded using a standard HD webcam placed above the middle of the screen. A single recording duration of 5 minutes has been shown to suffice for assessing stable spontaneous eye blink rate values (Barbato et al., 2000; Zaman & Doughty, 1997). Pupillometry data were collected using a RED-500 remote eye-tracking system (sampling rate (SR): 500 Hz; Sensomotoric Instruments, Teltow, Germany). Cardiovascular activity was measured using a BIOPAC MP 160 system (SR: 2000Hz; MP 160; Biopac systems, Inc). For cardiovascular recordings an ECG100C amplifier module with a gain of 2000, normal mode, 35 Hz low pass notch filter and 0.5 Hz/1.0 Hz high pass filter was included in the recording system. Disposable circular contact electrodes were attached according to the lead-II configuration. Isotonic paste (Biopac Gel 100) was used to ensure optimal signal transmission.

#### Physiological data analyses

##### Spontaneous eye blink rate

Spontaneous eye blink rate for each 5 minute recording was quantified by replaying the recorded videos and counting the single blinks manually by two separate evaluators. In case of a discrepancy >= 2 blinks this was repeated. The mean of both final evaluations (rounded to the lower integer) was used.

##### Cardiac data

Heart rate data (5 min recordings) were visually screened and manually corrected for major artifacts. We used custom MATLAB code to detect each R peak within the raw data. Specifically, the data was down sampled from 2000 to 200 Hz. Next, we applied the inverse maximum overlap discrete wavelet transform from MATLAB’s Wavelet Toolbox to reconstruct the heart rate data based on the 2^nd^ to the 4^th^ scale of the respective wavelet coefficients. The reconstructed data was squared and respective peaks were detected via MATLAB’s findpeaks function. The obtained peak locations were then subjected to a custom R script that used the RHRV toolbox for R (Rodríguez-Liñares et al., 2011) to compute heart rate and low frequency to high frequency ratio (LF/HF) of heart rate variability.

##### Pupil data

We used custom MATLAB code for pupil data analysis (5 min recordings). First, missing valus as a result of eye blinks or the like were removed. Next, pupil data were down-sampled from 500 Hz to 50 Hz. Outliers within a moving window of five seconds (250 data points) (mean ± 2 SD) were removed and linearly interpolated. The remaining data was smoothed using a robust weighted least squares local regression (‘rloess’ function within MATLAB’s Curve Fitting toolbox) with a span of 10 data points. Mean pupil size was then computed and averaged over both eyes for each assessment period.

##### Cue exposure

Participants underwent two cue exposure blocks on each testing day, the first before the temporal discounting task and the second before the sequential reinforcement learning task. Erotic pictures were selected via a google search. Neutral pictures were selected from the International Affective Picture system (IAPS; Lang, Bradley, & Cuthbert, 2008). Using MATLAB’s SHINE toolbox, images were converted to grayscale and matched according to luminance and contrast. In each exposure block, a total of 21 erotic or neutral photos were shown in randomized order twice in two separate rating rounds. During the first exposure round participants were instructed to rate each image on the arousal dimension. In a second exposure round they rated the pictures according to valence. To assess arousal, participants were asked how much arousal the image evoked. Arousal was rated on a visual-analogue scale presented on the screen ranging from “not arousing at all” to “very arousing”. After all images were rated based on arousal, the images were presented a second time in randomized order, for subsequent valence ratings. Here participants indicated the emotional valence evoked by the image on a visual analogue scale ranging from “very negative” to “very positive”. Each image was presented for at least 6 seconds until the participant responded. Between successive image presentations a fixation cross was shown for 3 seconds superimposed on a grey background screen with similar luminance as the images. Each cue exposure block lasted approximately 10 minutes.

#### Temporal discounting task

After completing the first cue exposure block, participants performed 128 trials of a temporal discounting task on each testing day. On each trial, participants selected between a “smaller sooner” (SS) reward available immediately and a “larger later” (LL) reward available after a particular delay in days. SS rewards were fixed to 20 € throughout the trials. LL magnitudes were computed by multiplying the SS reward with [1.01, 1.02, 1.05, 1.10, 1.15, 1.25, 1.35, 1.45, 1.65, 1.85, 2.05, 2.25, 2.65, 3.05, 3.45, 3.85] or [1.01, 1.03, 1.08, 1.12, 1.20, 1.30, 1.40, 1.50, 1.60, 1.80, 2.00, 2.20, 2.60, 3.00, 3.40, 3.8]. The temporal delays of the LL rewards ranged from 1 day to 120 days in eight steps [1, 3, 5, 8, 14, 30, 60,120] or [2, 4, 6, 9, 15, 32, 58, 119] days, respectively. LL magnitudes and delays were counterbalanced across the testing days and participants. As in previous studies (Green et al., 1997; Wagner et al., 2020) all choice options were hypothetical. Notably, discount rates for real and hypothetical rewards show a high correlation and similar neural underpinnings (Bickel et al., 2012).

#### Sequential reinforcement learning task

After completing the second cue exposure block, participants performed 300 trials of a modified version of the original two-step task by Daw et al. (2011). Based on suggestions by Kool et al. (2016) we modified the outcome stage by replacing the fluctuating reward probabilities (reward / no reward) with fluctuating reward magnitudes (Gaussian random walks with reflecting boundaries at 0 and 100, and standard deviation of 2.5). This task version has been successfully applied in a number of recent papers from our group (Mathar et al., 2022; Wagner et al., 2022).

In short, each trial comprised two successive decision stages. In the 1^st^ stage (S1), participants chose between two options represented by abstract geometrical shapes. Each S1 option probabilistically led to one of two 2^nd^ stage (S2) states that again comprised two choice options represented by abstract geometrical shapes. Which S2 stage state was presented depended probabilistically on the S1 choice according to a fixed common (70% of trials) and rare (30% of trials) transition scheme. The S2 stage choice options each led to a reward outcome. To achieve optimal performance, participants had to learn two aspects of the task. They had to learn the transition structure, that is, which S1 stimulus preferentially (70%) led to which pair of S2 stimuli. Further, they had to infer the fluctuating reward magnitudes associated with each S2 stimulus. We used different but matched task versions for the two testing days (erotic/neutral cue exposure, counterbalanced). Task versions used different S1 and S2 stimuli, and different S2 random walks (same rewards but with reversed order).

On each testing day, participants underwent extensive self-paced, computer-based instructions. Instructions provided detailed information about the task structure, the fixed transition probabilities between stages S1 and S2 and the fluctuating reward outcomes in S2. Participants were instructed to earn as much reward points as possible, and that following task completion reward points were converted to € such that they could win a bonus of up to 4€ in addition to their reimbursement of 10€/h. They performed 20 practice trials prior to task performance, with different random walks and stimuli.

Both tasks were implemented using the Psychophysics Toolbox Version 3 (PTB-3) running under MATLAB (The MathWorks ©).

#### Demographic and psychological screening

At the end of the first testing day, participants completed a brief survey on demographic data, socioeconomic status, weight and height (for calculating Body Mass Index (BMI)), trait impulsivity (Barratt-Impulsiveness Scale (BIS-15; German Version: Meule et al., 2020; Spinella, 2007), Behavioral Inhibition and Behavioral Activation System (BIS/BAS scale; German Version: Strobel et al., 2001; Carver & White, 1994), severity of depressive symptoms (Beck Depression Inventory-II (BDI-II; German Version: Kühner et al., 2007; Beck et al., 1996) and sleep duration (mean of the past 30 days). Please see *SI* (*table S1*) for related summary statistics.

#### Analysis of cue exposure on self-reported and physiological arousal

We used mixed effects regression models to assess the impact of erotic vs. neutral cue exposure on participants’ arousal ratings and changes compared to baseline in pupil dilation as well as heart rate and LF/HF heart rate variability ratio. For arousal ratings we modeled exposure round (before temporal discounting vs. before sequential reinforcement learning) and cue condition (erotic vs. neutral) as well as their interaction term as between subject effects, and round and cue also as within effects. The same mixed models were set up for testing cue exposure effects on changes relative to baseline in all three physiological markers of arousal.

#### Model-agnostic analysis of task performance

As a model-agnostic measure of temporal discounting performance, we examined fraction of LL choices, as well as mean RTs, as a function of cue exposure condition (erotic vs. neutral) with participants as a random effect using a (generalized) mixed effects regression approach. For the sequential RL task we used (generalized) mixed effects regression models to examine S1 stay/shift choice patterns, as well as S2 RTs. Stay probabilities in S1, were modeled as a function of previous reward, state transition (common vs. rare) and cue exposure condition. All three predictors as well as their interactions were modeled as between subject effects. They were also modeled as main effects on the within subject level. S2 RTs were modeled as a function of transition, cue exposure as well as their interaction on the between subject level with the inclusion of the main effects of both predictors on the within subject level. In line with our modeling analyses, task data were filtered such that trials with implausible fast or slow RTS (see details on DDM modeling) were excluded.

#### Computational modeling

##### Temporal discounting task

We applied a standard hyperbolic discounting model to describe how subjective value changes as a function of LL reward magnitude and delay:

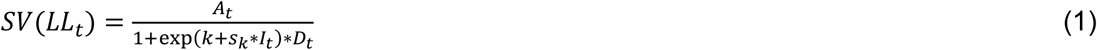

Here, *A_t_* is the reward magnitude of the LL option on trial *t*, *D_t_* is the LL delay in days on trial *t* and *I_t_* denotes the dummy-coded predictor of exposure condition. The model has two free parameters: *k* is the hyperbolic discounting rate (here modeled in log-space) and *s_k_* models a potential additive effect of erotic (vs. neutral) cue exposure on temporal discounting.

##### Temporal discounting task, softmax action selection

In a first modeling scheme, we used a softmax function to link subjective values of LL and SS rewards in each trial with participants’ choices:

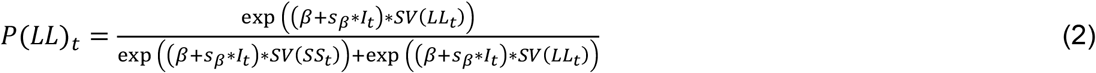

Here, *SV* is the subjective value of the larger but later reward according to Eq. 1 and *β* is the inverse temperature parameter, modeling choice stochasticity. *SV*(*SS_t_*) was fixed at 20 and *I_t_* is again the dummy-coded predictor of supplementation condition, and *s_β_* models a potential modulatory effect of erotic cue exposure on *β*. This softmax model was used as a reference model for comparison purposes (*SI* - posterior predictive checks, figure S1) for a drift diffusion model implementation delineated below.

##### Temporal discounting task, drift diffusion model (DDM) implementation

In a next step, we replaced softmax action selection with a series of drift diffusion model (DDM)-based choice rules. The DDM belongs to the family of sequential sampling models. In these models, binary decisions are assumed to arise from a noisy evidence accumulation process that terminates as soon as the evidence exceeds one of two response boundaries. In all DDM implementations, the upper boundary was defined as the selection of the LL option, whereas the lower boundary was defined as choosing the SS option. RTs for choices of the SS option were multiplied by −1 prior to model fitting. Prior to modeling, we filtered the choice data using percentile-based RT cut-offs, such that on a group-level the fastest and slowest one percent of all trials according to RTs were excluded from modeling, and on an individual subject level the fastest and slowest 2.5 percent were further discarded. With this we avoid that outlier trials with implausible short or long RTs bias the results. Further, we decided to exclude participants from data analysis with more than 20% detected outlier trials as indicative of improper task engagement. In the temporal discounting task this was the case for one participant.

We first implemented a null model (DDM0) without any value modulation of the drift-rate *v*:

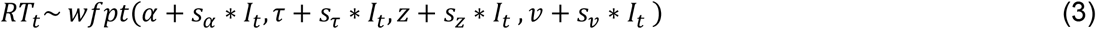

The starting point bias *z* was fitted to the data, such that z>.5 reflected a bias towards the LL boundary, and z<.5 reflected a bias towards the SS boundary. As in previous work (Fontanesi et al., 2019; Pedersen et al., 2017; Peters & D’Esposito, 2020), we then set up temporal discounting DDMs with a modulation of drift-rates by the difference in subjective values between the LL and SS options. First, we set up an implementation with a linear modulation of drift-rates (DDM_lin_) (Pedersen et al., 2017):

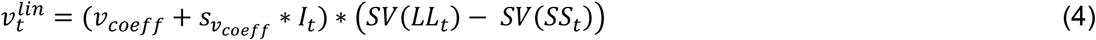

We next examined a DDM with non-linear (sigmoid) trial-wise drift-rate scaling (DDM_S_) that has recently been reported to account for the value-dependency of RTs better than the DDM_lin_ (Fontanesi et al., 2019; Peters & D’Esposito, 2020; Wagner et al., 2020, 2022). In this model, the scaled value difference from Eq. 17 is additionally passed through a sigmoid function with asymptote v_max_:

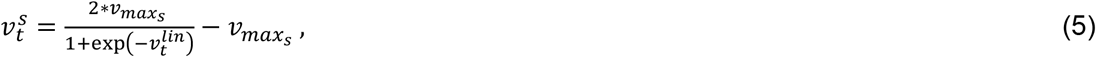

with

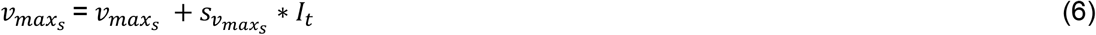

All parameters were again allowed to vary according to the cue exposure condition, such that we included *s_x_* parameters for each parameter *x* that were multiplied with the dummy-coded condition predictor *I_t_* to model potential effects of erotic cue exposure on model parameters.

#### Sequential RL task

As in our previous work (Wagner et al., 2022), we implemented a slightly modified version of the original hybrid reinforcement learning model for the seq. reinforcement learning task (Daw et al., 2011; Otto et al., 2015) to analyze model-free and model-based contributions to behavior. As in Otto et al., (2015) the model contains separate coefficients to model contributions of model-based and model-free values of S1 choices, instead of a single weighting parameter (Daw et al., 2011). In addition, our model includes a forgetting rate parameter of unchosen option values, as suggested previously (Toyama et al., 2017, 2019). Note that the original model without a forgetting process is contained as a nested version in this extended model (i.e. in the case of forgetting rate of zero). The model updates model-free state-action values (*Q_MF_*-values, Eq. 7, 8) in both stages S1 & S2 via a prediction error scheme (Eq. 9, 10). In S1, model-based state-action values (*Q_MB_*) are then computed from the transition and reward estimates using the Bellman Equation (Eq. 11). To account for potential modulatory effects of erotic vs. neutral cue exposure, we included additive ‘shift’ parameters *s_x_* for each parameter *x* that were multiplied by dummy-coded exposure condition predictors *I_t_* (= 1, erotic; = 0, neutral).

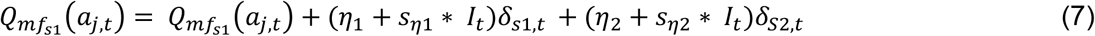

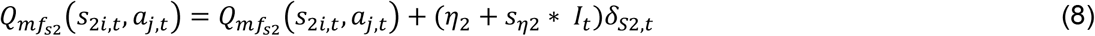

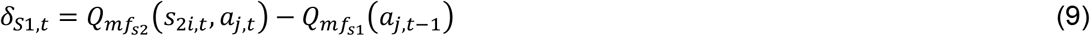

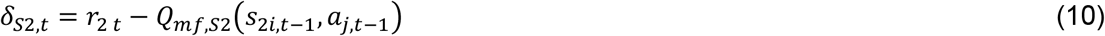

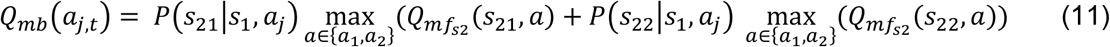

Here, *i* indexes the two different second stage states (*S*_21_, *S*_22_), *j* indexes actions *a* (*a*_1_, *a*_2_) and *t* indexes the trials. Further, *η*_1_ and *η*_2_ denote the learning rate for S1 and S2, respectively. S2 model-free *Q*-values are updated by means of reward (*r*_2,*t*_) prediction errors (*δ*_*S*2,*t*_) (Eq. 2, 4). To model S1 model-free *Q*-values we allow for reward prediction errors at the 2nd-stage (Eq. 4) to influence 1st-stage *Q*-values (Eq. 1). In addition, *Q*-values of all unchosen stimuli were assumed to decay with forgetting rate *γ_s_* (Toyama et al., 2017, 2019) towards the mean of the reward outcomes (0.5) according to Eq. 6 and 7:

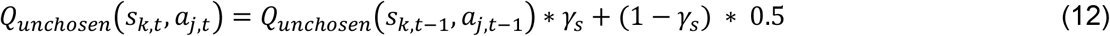

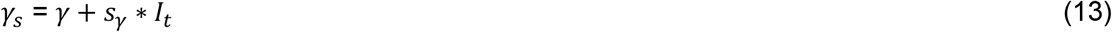

with *k ϵ* {1, 21, 22} indexing the first (S1) and the two second stage (S21, S22) states.

This learning model was then combined with two different choice rules: softmax action selection, and the drift diffusion model (Shahar et al. 2019; Wagner et al., 2022).

##### Sequential RL task, softmax action selection

We first implemented a standard softmax action selection scheme to link learned Q-values with participants’ choices. Softmax action selection models choice probabilities as a sigmoid function of value differences (Sutton & Barto, 1998). In this regard, S1 choice probabilities are modelled via weighting of S1 model-free and model-based *Q*-values through a softmax function. Similarly, S2 stage action selection is modelled as a function of weighted model-free *Q*-values (Eq. 14, 15). An additional parameter *ρ* was included to model 1st-stage choice perseveration, *rep*(*a*) that is set to 1 if the previous S1 choice was the same and is zero otherwise.

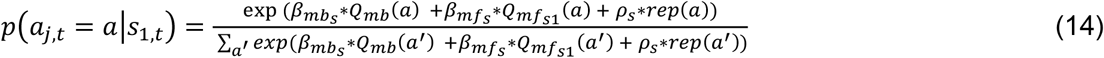

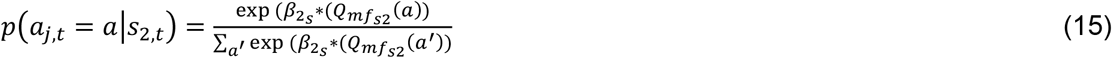

with:

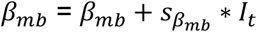

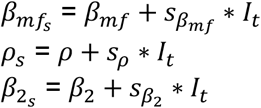

To account for potential modulatory effects of erotic vs. neutral cue exposure, we included additive ‘shift’ parameters *s_x_* for each parameter *x* that were multiplied by dummy-coded supplementation predictors *I_t_* (= 1, erotic; = 0, neutral). Note, that this softmax model was used for comparison purposes (*SI – posterior predictive checks*, figure S1) for a more advanced drift diffusion model framework delineated below.

##### Sequential RL task, drift diffusion model (DDM) implementation

Similar to modeling of the temporal discounting task, and as reported in Wagner et al. (2022) we replaced the standard softmax action selection with a series of DDM-based choice rules to more comprehensively examine modulatory effects of erotic (vs. neutral) cue exposure on choice dynamics. For each stage S1/S2 of the seq. RL task, the upper boundary was defined as selection of one stimulus, whereas the lower boundary was defined as selection of the alternative stimulus. RTs for choices of the alternative option were multiplied by −1 prior to model fitting. We used a percentile-based cut-off for S1 and S2 RTs similar to the one described in the temporal discounting DDM section. Based on this cut-off, three participants had fast/slow outlier responses in more than 20% of the trials and were thus disregarded from data analysis.

We then first examined a null model (DDM_0_) without any value modulation. Trial-wise RTs on each stage S1 & S2 are assumed to be distributed according to the Wiener-First-Passage-Time (*wfpt*):

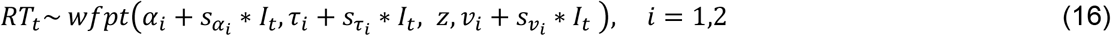

Here, boundary separation parameters *α_i_* model the amount of evidence required before committing to a decision in each stage *S_i_*, non-decision time *τ*_*i*_ model components of the RT that are not directly implicated in the choice process, such as motor preparation and stimulus processing. The starting point bias *z* models a bias towards one of the response boundaries before the evidence accumulation process starts. This was set to .5 for both stages, as the boundaries were randomly associated with the choice options on each stage. The drift-rate parameters *vi* model the speed of evidence accumulation. Note that for each parameter *x*, we also included a parameter *s_x_* that models potential modulatory effects of erotic (vs. neutral) cue exposure (coded via the dummy-coded condition predictor *I_t_*).

As in previous work (Fontanesi et al., 2019; Pedersen et al., 2017; Peters & D’Esposito, 2020), we then set up hybrid RL DDMs with modulation of drift-rates by value differences between the respective choice options, separately for each stage. First, we set up a model with a linear modulation of drift-rates (DDM_lin_) (Pedersen et al., 2017). For S1, this yields

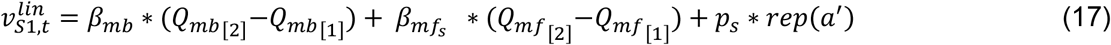

and the drift-rate in S2 is computed as

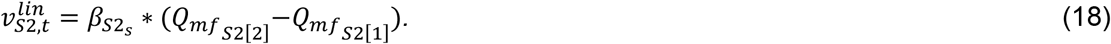

with

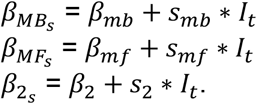

We next set up a DDM with non-linear (sigmoid) drift-rate modulation (DDM_S_) that has recently been shown to better account for the value-dependency of RTs compared with the DDM_lin_ (Bartels et al., 2018; Fontanesi et al., 2019; Peters & D’Esposito, 2020; Wagner et al., 2020, 2022). In this model, the scaled value differences from Eq. 17 & 18 are additionally modulated by a sigmoid function with asymptote 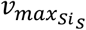

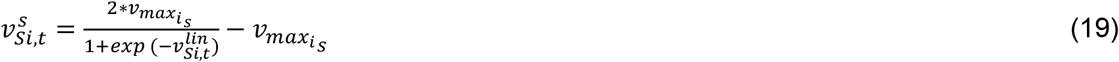

with

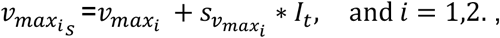

#### Hierarchical Bayesian model estimation

Models were fit to all trials from all participants with less than 20% outlier trials (fast/slow trials based on percentile-related cut-offs, see above) using a hierarchical Bayesian modeling approach with separate group-level distributions for all parameters of the neutral (baseline) condition and additional shift parameters *s_x_* to model erotic cue exposure effects on all parameters. Model fitting was performed using MCMC sampling as implemented in Stan (Stan Development Team, 2020) running under R (Version 3.5.1) and the RStan package (Version 2.21.0). For baseline group-level means, we used uniform and normal priors defined over numerically plausible parameter ranges (see code and data availability section for details). For all *s_x_* parameters modeling erotic cue exposure related effects on model parameters, we used normal priors with means of 0 and numerically plausible standard deviations (range=1-10). For group-level standard deviations we used half-cauchy distributed priors with location=0 and scale=2.5 for the temporal discounting data and uniform priors within plausible ranges (mean=0, range standard deviation=10-50) for the sequential reinforcement task data??. Sampling was performed with four chains, each chain running for 4000 (6000 for the seq. RL data) iterations without thinning after a warmup period of 3000 (5000 for the seq. RL data) iterations. Chain convergence was then assessed via the Gelman-Rubinstein convergence diagnostic 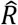 with 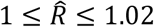 as acceptable values. For both tasks, relative model comparison was performed via the *loo*-package in R (Version 2.4.1) using the Widely-Applicable Information Criterion (WAIC) and the estimated log pointwise predictive density (elpd) which estimates the leave-one-out cross-validation predictive accuracy of the model (Vehtari et al., 2017). We then report posterior group distributions for all parameters of interest as well as their 80% and 95% highest density intervals (HDI). For erotic cue exposure effects, we report Bayes Factors for directional effects of parameter distributions of *s_x_*, estimated via kernel density estimation using R via the RStudio (Version 1.3) interface. These are computed as the ratio of the integral of the posterior difference distribution from 0 to +∞ vs. the integral from 0 to −∞. Using common criteria (Beard et al., 2016), we considered Bayes Factors between 1 and 3 as anecdotal evidence, Bayes Factors above 3 as moderate evidence and Bayes Factors above 10 as strong evidence. Bayes Factors above 30 and 100 were considered as very strong and extreme evidence respectively, whereas the inverse of these reflect evidence in favor of the opposite hypothesis.

#### Posterior Predictive checks

We carried out posterior predictive checks (*SI - posterior predictive checks*, figure S1, 2) to examine whether models could reproduce key patterns in the data, in particular the value-dependency of RTs (Peters & D’Esposito, 2020) and that of participant’s choices. For the seq. RL task, we extracted 500 unique combinations of posterior parameter estimates from the respective posterior distributions and used these to simulate 500 datasets using the Rwiener package (Version 1.3.3). We then show median RTs of observed data and the median RTs from the 500 simulated datasets for all DDMs as a function of value differences. Value differences for S1 were computed as the absolute difference between the maximum 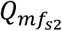 values of each S2 stage weighted by their respective transition probability. Value differences in S2 were computed as the difference in the actual reward values of the respective choice options. Similarly, we show that our models capture the dependency of participants’ stay probabilities in S1 on S1 value differences, and the dependency of their fraction of optimal (max[reward]) choices in S2 on S2 reward differences. For the intertemporal choice task, we binned trials of each individual participant into five bins, according to the absolute difference in subjective LL vs. SS (“decision conflict”), computed according to each participant’s median posterior *k* parameter from the DDM_S_ separately for the neutral and erotic cue exposure condition. For each participant and condition, we then plotted the mean observed RTs and the percentage of LL choices as a function of decision conflict, as well as the mean RTs and fraction of LL choices across 500 data sets simulated from the posterior distributions of the DDM_0_, DDM_lin_ and DDM_S_ and softmax model (choices only).

## Results

### Self-reported and physiological arousal

On each testing day, participants underwent a 5 minute baseline assessment of physiological arousal at rest and two (approx. 10 minutes) assessments of physiological and self-reported arousal during cue exposure (carried out before each decision-making task). First, measures of physiological arousal showed good test-retest reliability (intra class correlation coefficient (icc)) throughout the two baseline assessments except for LF/HF ratio of heart rate variability that only exhibited moderate reliability (table 1, figure 1b,c). Self-reported arousal during erotic compared with neutral cue exposure showed excellent test-retest reliability throughout the two exposure assessments (icc(Δarousal)=.94, p=4.81*10^−20); figure 1a).

**Table 1.**
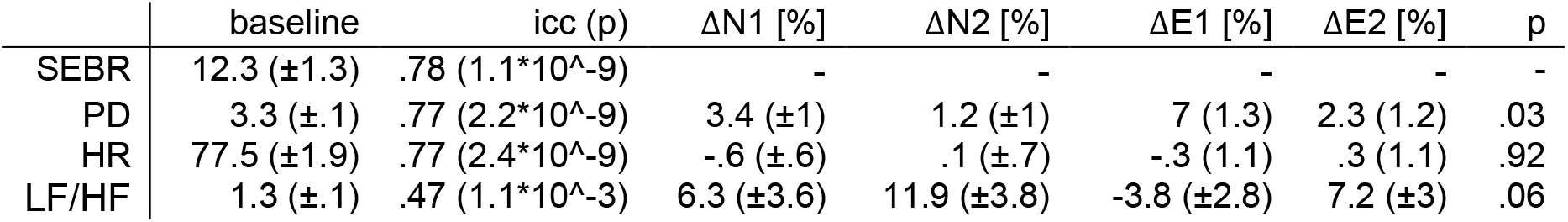
Mean (standard error) of physiological arousal markers during the two baseline sessions, test-retest reliability (intra class correlation coefficient (icc), p-value), and related percent change thereof during cue exposure (N=neutral, E=erotic; 1 = exposure round before temporal discounting, 2 = before reinforcement learning). The last column depicts p-values of the main effect of cue from mixed effects regression analysis. SEBR: spontaneous eye blink rate per minute (only assessed at baseline); PD: mean pupil dilation size; HR: heart rate; LF/HF: low frequency to high frequency ratio of heart rate variability. Sample size: N=39 (per measurement session).

**Figure 1.**
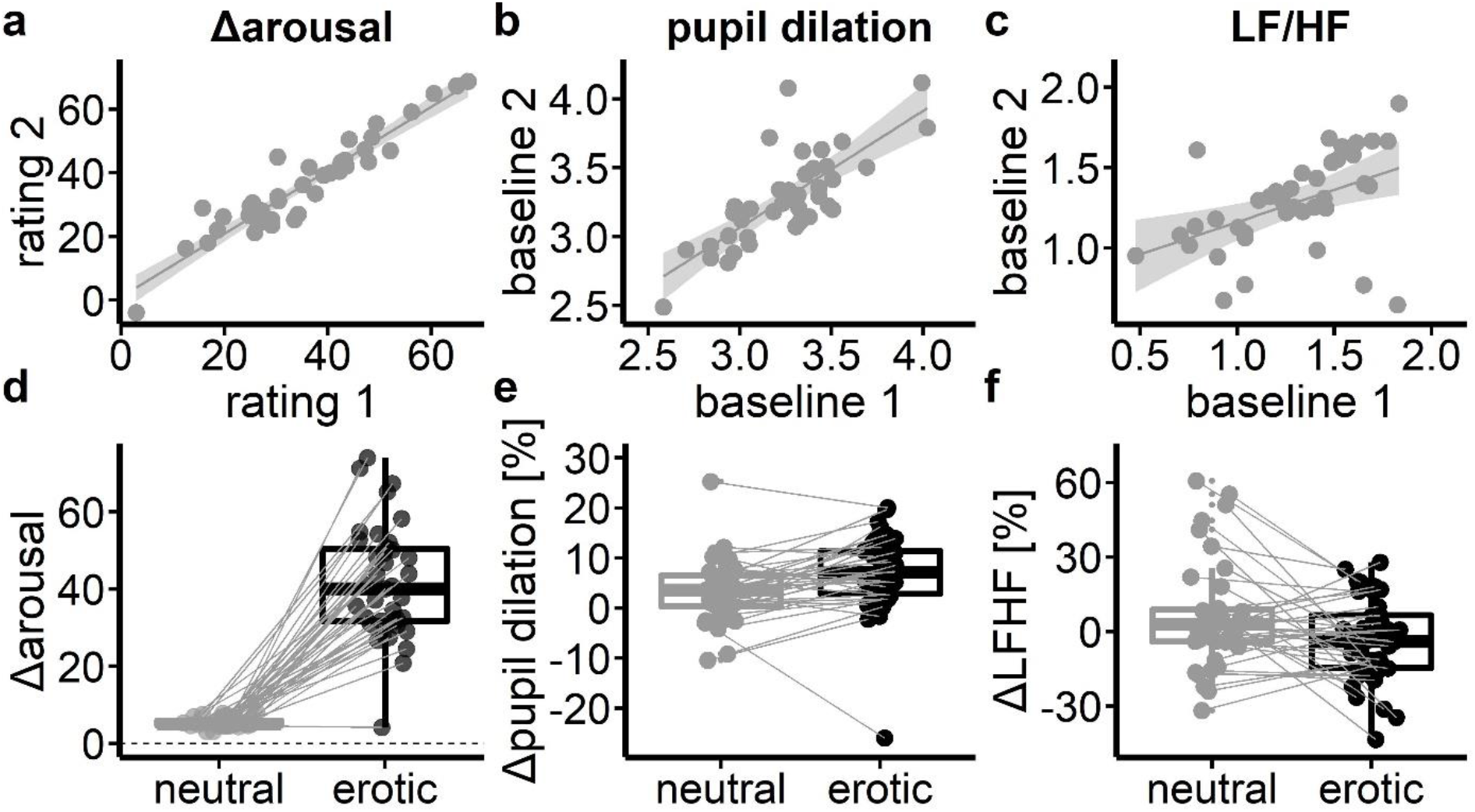
(a-c) Self-reported arousal (erotic – neutral) and pupil dilation, showed good to excellent test-retest reliability throughout the separate ratings (self-reported) and baseline assessments (autonomic). (d) Self-reported arousal was significantly increased during erotic as compared to neutral cue exposure and showed excellent reliability throughout the two exposure rounds. (e) Pupil dilation change in relation to baseline was higher during erotic compared to neutral cue exposure. (f) Low to high frequency ratio of heart rate variability was nominally reduced in relation to baseline during erotic vs. neutral cue exposure.

Next, we utilized mixed effects regression models to assess effects of erotic compared with neutral cue exposure on self-reported and autonomic markers of arousal. Erotic compared with neutral cue exposure was associated with increased self-reported arousal as revealed by a significant main effect of cue (*β*=17.91, *t*=15.0, p<2*10^−16^; figure 1d). Exposure round (1/2) was also a significant positive predictor of reported arousal (*β*=.46, *t*=2.18, p=.03), as well as the interaction of cue*round (*β*=.22, t=2.4, p=.02) implying that self-reported arousal even increased more strongly during erotic compared with neutral cue exposure between the first and the second exposure round. On the autonomic level we observed heightened pupil dilation during erotic vs. neutral cue exposure in relation to pupil dilation at baseline according to a significant main effect of cue (*β*=1.17, t=2.22, p=.03; figure 1e). In contrast to a positive effect of erotic cue exposure, pupil dilation decreased over the successive exposure rounds in general (*β*=−1.75, t=−6.67, p=6.86*10^−8) and this effect appeared to be even stronger for erotic vs. neutral exposure rounds (*β*=−.63, t=−3.24, p=.002). Post-hoc t-tests revealed that in relation to baseline, pupil size was increased in the first exposure round (t=3.14, p=3.29*10^−3), but not in the second exposure round during erotic vs neutral cue exposure (t=0.98, p=0.3325). Heart rate showed no modulation neither by cue (*β*=.04, t=.1, p=.92) nor by exposure round (*β*=.34, t=1.41, p=.17)). LF/HF ratio of heart rate variability was nominally decreased during erotic vs. neutral cue exposure in relation to baseline assessment (*β*=−3.59, t=−1.89, p=.06; figure 1f) indicating a marginally stronger parasympathetic control of heart muscle activity. LF/HF ratio was also modulated by exposure round (*β*=4.27, t=4.07, p=.0002) and we observed a nominal interaction of cue and exposure round (*β*=1,44, t=1.81, p=.08). Similar to the changes observed in pupil dilation, post-hoc t-test revealed a significant difference in LF/HF ratio between erotic and neutral cue exposure in the first exposure round (t=−2.62, p=.01) but not in the second (t=−.98, p=.33) in relation to baseline LF/HF ratio.

To reveal if subjective arousal corresponded with autonomic measures, we tested whether self-reported arousal during erotic vs neutral cue exposure would predict changes in autonomic arousal markers, that is changes in pupil dilation and LF/HF heart rate variability ratio. While cue exposure related increases in pupil dilation were not predicted by self-reported arousal (erotic vs. neutral) (r=−.13, t=−0.78, p=0.44), reductions in participants’ LF/HF ratio following the first exposure were stronger with greater self-reported arousal (r=−.35, t=−2.26, p=0.03, figure 2b). Spontaneous eye blink rate is discussed as a proxy measure of central catecholamine transmission (Jongkees & Colzato, 2016), and Van Slooten et al. (2017) showed that participants with lower eye blink rate showed stronger reward expectation and surprise related pupil dilation changes than participants with higher eye blink rate. Thus, we tested whether individual eye blink rates would predict erotic cue exposure related changes in pupil dilation in an exploratory attempt. In accordance with the findings from Van Slooten et al. (2017) we found that individuals with lower eye blink rate at rest showed stronger increases in pupil dilation during erotic cue exposure (robust regression: R^2^=.13, t=−2.22, p=0.03; figure 2a).

**Figure 2.**
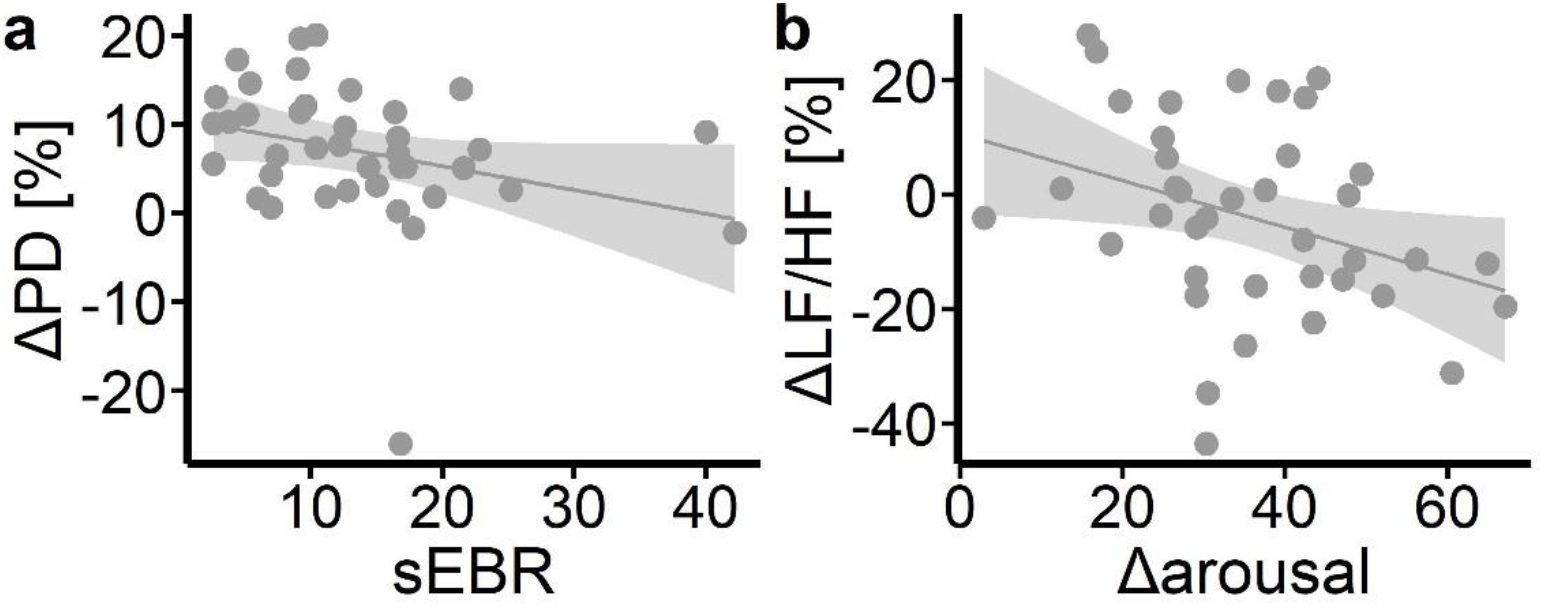
(a) Spontaneous eye blink rate (sEBR) at baseline predicted % change in pupil dilation (ΔPD) following erotic cue exposure. (b) Self-reported arousal was associated with change in low frequency to high frequency heart rate variability ratio.

### Temporal discounting task

#### Model-agnostic analysis

According to a mixed effects regression with cue as fixed and participants as random effects on individual fraction of LL choices, erotic compared with neutral cue exposure increased temporal discounting on a nominal level (%LL choices, erotic (neutral): 52.12 (54.4) ± 3.74 (3.34); t=−1.9, p=.07; figure 3a,b). We also computed a mixed effects regression model that included delay and LL magnitude (as median splits) as additional predictors both on the fixed and random level in the model. Here, we observed a significant modulation of the tendency to choose the LL option by both delay (*β*=−1.42, z=−10.75, p<2*10^−16) and LL magnitude (*β*=2.23, z=12.36, p<2*10^−16), and their interaction (*β*=−.47, z=−3.94, p=5.52*10^−5). Similar to the simpler mixed model, we found a marginal significant negative effect of cue condition (erotic vs. neutral) on LL choice tendency (*β*=−.07, z=−1.95, p=.05). In addition, all three interactions of cue condition with delay length (*β*=0.1, z=2.79, p=0.005), LL magnitude (*β*=0.11, z=3.19, p=0.001) and their three-way interaction (*β*=0.09, z=2.46, p=0.01) showed significant effects regarding participants’ proportion of LL choices. We found no evidence for a modulation of participants’ response times by erotic vs. neutral cue exposure (*β*=−.02, t=−.28, p=0.78; figure 3c,d).

**Figure 3.**
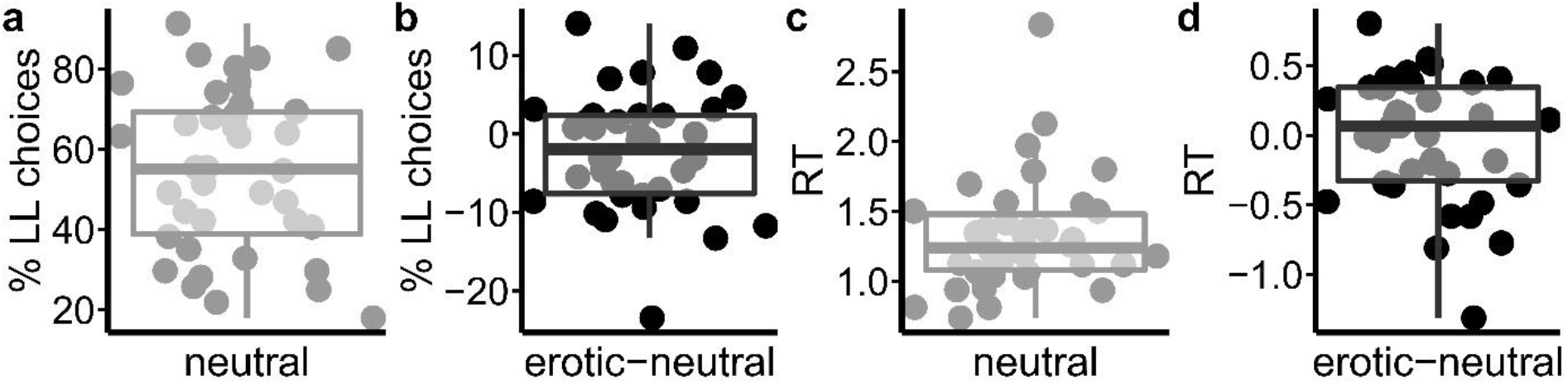
(a) Proportion of LL choices following the neutral cue exposure in the temporal discounting task. (b) Difference of % LL choices following erotic compared with neutral cue exposure. (c) Median RTs following the neutral cue exposure and (d) difference of RTs following erotic vs. neutral cues.

#### Drift diffusion model (DDM)

First, we examined the model fit (WAIC, elpd) of three implementations of the DDM that varied in the way they accounted for the modulation of trial-wise drift rates by value differences. A DDM with nonlinear drift-rate scaling (DDM_s_) (Fontanesi et al., 2019; Peters & D’Esposito, 2020; Wagner et al., 2020, 2022) accounted for the temporal discounting data best when compared to a DDM with linear scaling (DDM_lin_) (Pedersen et al., 2017) and a null model without value modulation (DDM_0_) (Table 2). We also compared the three DDMs to a softmax model with respect to the proportion of correctly predicted binary (SS vs. LL) choices. The DDMs predicted participants’ choices numerically on par with the softmax model, whereas the DDM_lin_ and even more so the DDM_0_ performed substantially worse (see SI table S1). Posterior predictive checks of the best-fitting model (DDM_s_) revealed that it accurately reproduced the effect of value differences (decision conflict) on participants’ RTs and the proportion of LL choices (*SI* figure S1). Note that we previously reported extensive parameter recovery analyses for this model (Peters & D’Esposito, 2020; Wagner et al., 2020). We then examined participants’ discounting behavior in greater detail via the posterior distributions of the group-level means of the DDMs parameters (figure 3).

**Table 2.**
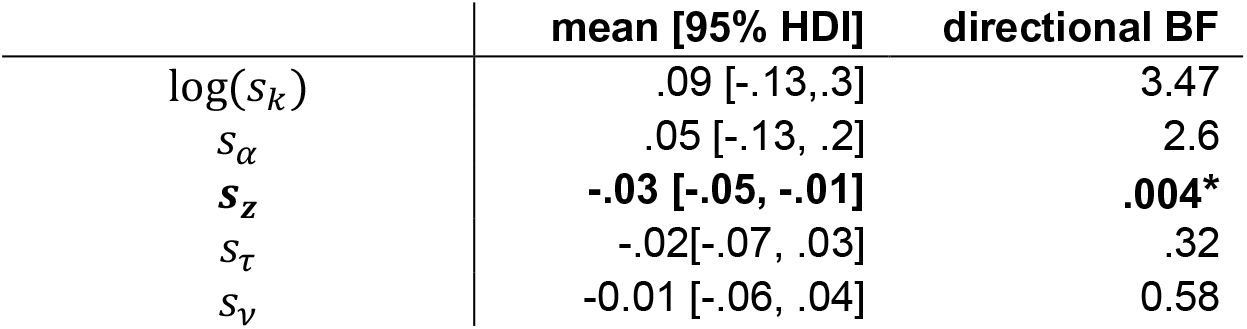
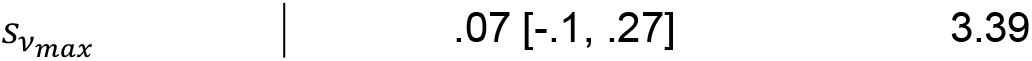
Erotic cue exposure related changes in group-level parameter means of the DDM_s_ of the temporal discounting data. We report mean and 95% HDIs for the ‘shift’ parameters and Bayes Factors for directional effects (erotic>neutral). * Denotes strong evidence (dBF<.1) for a negative effect of erotic cue exposure on the shift in the starting point bias parameter ***s_z_***.

We found no evidence for a general bias towards the SS or the LL option following the neutral cue exposure, as the 95% HDI for the starting-point bias parameter *z* overlapped with 0.5 (figure 4b). Furthermore, trial-wise drift rates increased with increasing value differences, such that the 95% HDI for the drift-rate coefficient parameter *ν* did not include 0 (see figure 4 h). Mirroring the tendency to choose the LL option less often following erotic vs. neutral cue exposure, we observed a negative shift in the starting point bias towards the SS option (*s*_z_; Figure 4c & Table 2). The stronger the individual shift in starting point bias was following erotic cue exposure, the more often participants chose the SS option compared with their choices following neutral cue exposure (r=.44, p=.005). All other parameters were not modulated by erotic cue exposure (95% (& 80%) HDI’s of all other shift parameters modeling condition effects overlapped with zero, figure 3).

**Figure 4.**
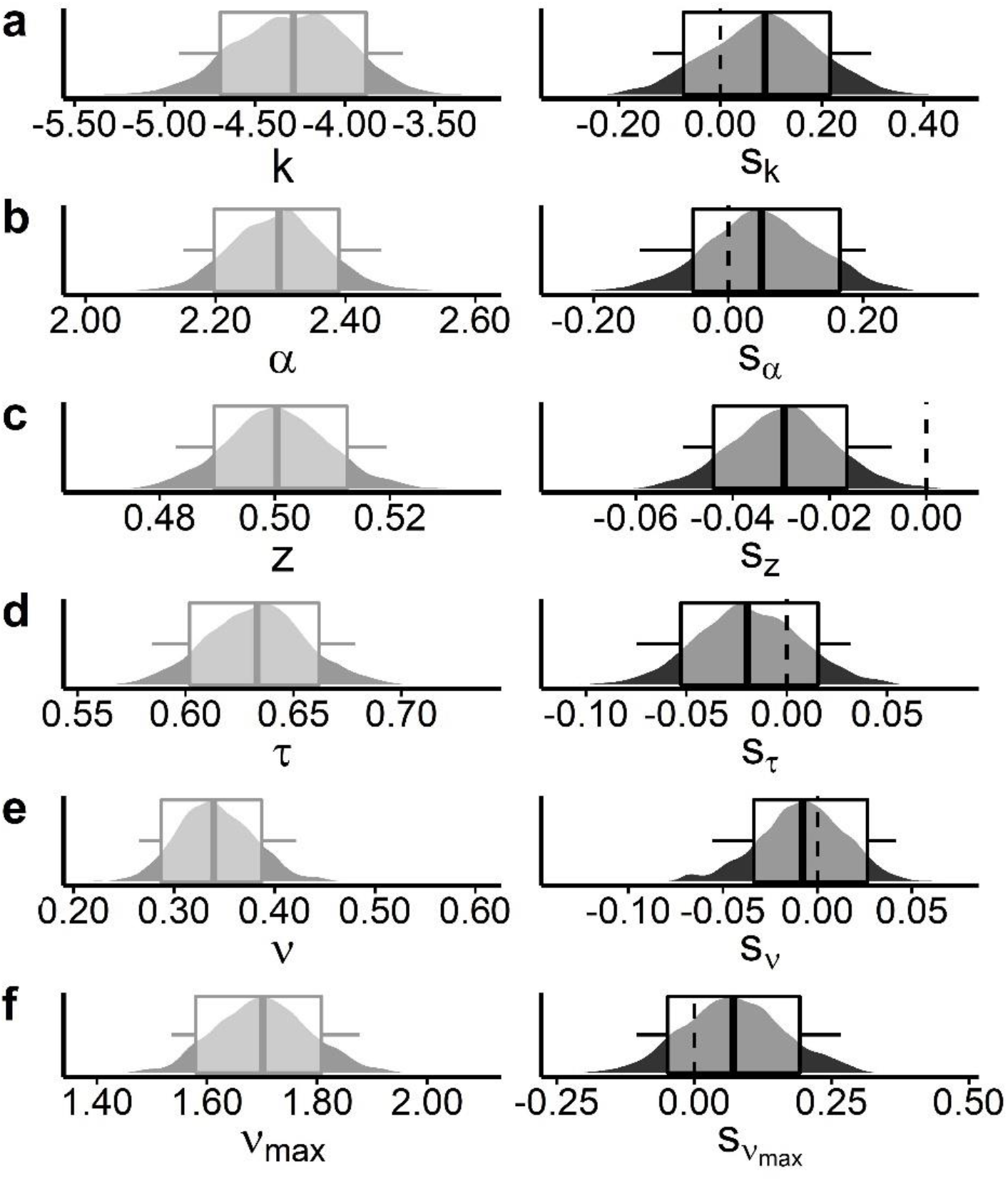
Posterior parameter distributions of the DDM_s_ group-level means of the temporal discounting data following neutral cue exposure (left column, grey plots) and their respective shifts as changes related to erotic vs. neutral cue exposure (right column, black plots). Boxplots depict 80% and 95% HDIs. Depicted parameters are (a) discount rate log(*k*) and shift *s_k_*, (b) decision-threshold *α* with shift *s_α_*, (c) starting point bias *z* and respective shift *s_z_*, (d) non-decision time *τ* with shift *s_τ_*, (e) drift-rate *ν* and related shift *s*_*ν*_, and (f) drift rate asymptote *v_max_* with shift 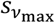. Only (c) starting point bias *z* showed strong evidence for a negative shift following erotic cue exposure (s_z_, 95% HDI < 0), such that participant’s starting point (bias) shifted towards smaller-sooner options.

#### Sequential reinforcement learning task

##### Model-agnostic analyses

Since recently the reliability of seq. reinforcement learning performance has been called into question (Enkavi et al., 2019), we assessed whether this holds true for the slightly modified task version we used here. We tested test-retest reliability between exposure sessions of the individual beta weights from the reward*transition interaction of the mixed effects regression model as an index of model-based control. Despite the cue exposure manipulation, test-retest reliability appeared to be good with considerably higher icc values (icc=0.65, p=6.29*10^−6) than reported in Enkavi et al. (2019).

Next, we tested for effects of erotic cue exposure on task performance. Exposure to erotic compared with neutral cues had no significant effect on participants’ average payout per trial (erotic: 61.94 (± .54) vs neutral: 61.53 ± .76 [mean (±SE)]; t=−.82, p=.42).

We then used a generalized mixed effects regression approach to examine participants’ stay/shift behavior in the first decision stage S1 as a function of previous reward receipt, state transition (common vs. rare), cue exposure and their interactions. We observed a main effect of reward (*β*=.14, SE=.03, p=7.14*10^−8; table 2), reflecting a model-free contribution to behavior, and a reward*transition interaction indicating that participants also incorporated a model-based reinforcement learning strategy (*β*=.43, SE=.05, p=2.29*10^−18; table 2, figure 5a). We observed a significant negative main effect of cue exposure (erotic vs. neutral) on participants’ probability to choose the same S1 option as in the preceding trial (*β*=−.11, SE=.02, p=4.52*10^−8; figure 5b), as well as a significant interaction of reward*transition*cue exposure on participants’ stay probabilities (*β*=−.07, SE=.02, p=1.72*10^−3; figure 5b). Thus, erotic cue exposure decreased participants’ tendency to choose the same S1 option as in the preceding trial irrespective of the reward received and the transition experienced. More specifically, erotic cues reduced signatures of model-based control in participants’ stay probability patterns of 1^st^ stage choices. Post-hoc we tested which specific transition*reward combination was affected. Here, we found that following rare transitions and lower than expected rewards (i.e. smaller than the mean of the last 20 trials) participants were less likely to choose the same S1 option again (Δp(stay)=−.07, t=−4.4, p=9.63*10^−5; figure 5b). We also found a reduction in the probability to choose the same option in S1 following a common transition in combination with a greater than expected reward (Δp(stay)=−.03, t=−2.31, p=0.03; figure 5b), although this fell short of surviving correction for multiple comparisons (False Discovery Rate)

**Figure 5.**
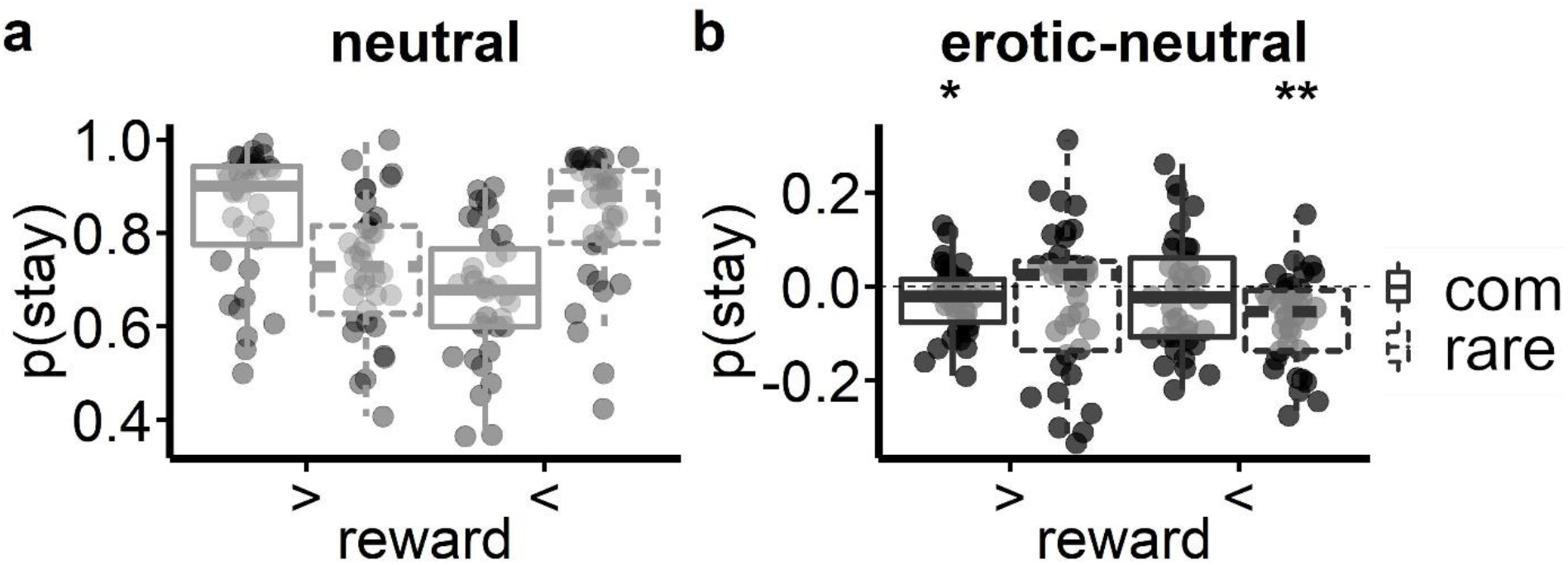
(a) Participants’ mean probabilities to choose the same S1 option again as in the preceding trial (p(stay)) dependent on reward height (‘>’ (‘<’) greater or equal (smaller) than the mean of the last 20 trials) and transition type (common, rare) following (a) neutral cue exposure and (b) respective differences following exposure to erotic cues. * (**) denote p-values <.05 (.001) of post-hoc t-tests.

In a similar fashion we examined potential effects of erotic cue exposure on participants’ S1 and S2 RTs. We first tested for a general effect of cue exposure on trial-wise RTs in the first decision stage S1 in a mixed effects regression model including the same predictors as the model for predicting S1 stay probabilities (see above). Here, we found no significant main effects or interactions for S1 RTs. There was only a nominal effect of preceding transition type (β=3.42*10^−3, SE=2.13*10^−3, p=.09) and cue exposure (β=3.03*10^−3., SE=1.69*10^−3, p=.07) on S1 RTs indicating that participants tended to respond slightly slower following a rare transition in the preceding trial and subsequent to erotic vs. neutral cue exposure in general. There was also only a nominal interaction of reward*transition*cue exposure (β=−2.98*10^−3, SE=1.81*10^−3, p=.09) hinting at slight reductions in the rare transition related slowing of S1 RTs following erotic vs. neutral cue exposure.

Increased S2 RTs following rare transitions are another indicator of model-based control similar to the above reported reward*transition interaction in S1 choice probabilities (Otto et al., 2015; Shahar et al., 2019; Wagner et al., 2022). Thus, next we computed a mixed effects regression model with transition, cue type and their interaction as fixed effects on S2 RTs. As previously reported (Otto et al., 2013; Shahar et al., 2019; Wagner et al., 2022), we observed a significant slowing of S2 RTs (β=.08, SE=8.5*10^−3, p=1.3*10^−11), figure 6a) following rare vs. common transitions. We also found a significant slowing of S2 RTs following erotic vs. neutral cue exposure (β=.01, SE=6.21*10^−3, p=.04, figure 6a). In contrast to the evidence of erotic-cue exposure associated attenuation of model-based control in the stay probability analysis, we did not find a significant transition*cue exposure interaction effect on S2 RTs (β=−8.55*10^−4, SE=6.22*10^−3, p=.89).

**Figure 6.**
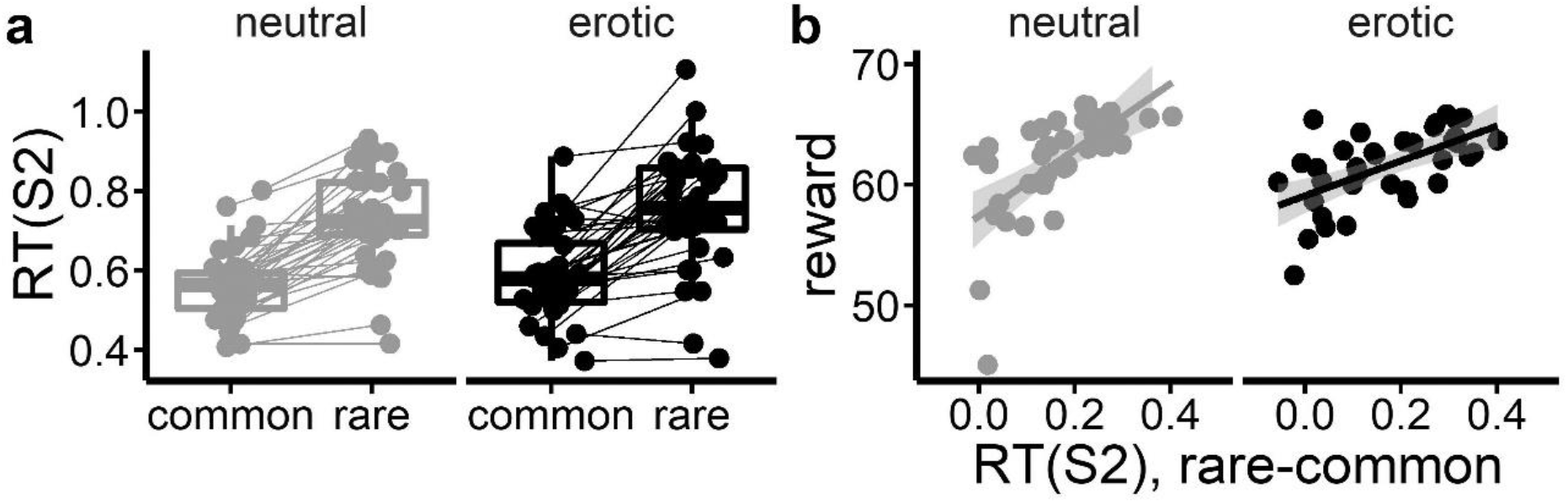
(a) Participants’ median RTs in stage S2 as a function of encountered transition following neutral and exposure to erotic cues. (b) Correlation of participants’ mean rewards per trial and the differences in median S2 RTs following a rare vs. a common state transition following neutral and erotic cue exposure.

In the modified task version that we employed, and in contrast to the original sequential reinforcement learning task from Daw et al., (2011), increased model-based control actually pays off as it leads to increased payout (Kool et al., 2016). This was reflected in a significant association of participants’ mean payoffs per trial and RT differences in stage S2 following a rare vs. a common transition (neutral (erotic): r=.64(.60), p=2.69*10^−5(1.15*10^−4); figure 6b).

##### Drift diffusion model (DDM)

Next, we implemented a DDM choice rule to make use of both choices and RT distributions (Shahar et al., 2019, Wagner et al., 2021, 2022). First, we examined the model fit (WAIC, elpd) of three implementations of the DDM that varied in the way they accounted for the modulation of trial-wise S1 & S2 drift-rates by Q-value differences. The DDM with nonlinear drift-rate scaling (DDM_s_) (Fontanesi et al., 2019; Peters & D’Esposito, 2020; Wagner et al., 2022) accounted for the data best when compared to a DDM with linear scaling (DDM_lin_) (Pedersen et al., 2017) and a null model without value modulation (DDM_0_) (see *SI* - table 3). Note, that we also compared the three DDM_s_ with a softmax model formulation according to the proportion of correctly predicted binary choices in S1 and S2, respectively. Since the softmax model only accounts for choice data, it shows the highest predictive accuracy. However, the DDM_s_ predicted participants’ choices better than the DDM_lin_ and the DDM_0_ (see *SI - table S3*). Importantly, posterior predictive checks for the best-fitting model DDM_s_ showed that it reproduced the impact of decision conflict on participants’ RTs and choice patterns (see *SI* - *figure S2*).

**Table 3.** Erotic cue exposure associated changes in group-level means of the DDM_s_ for the sequential reinforcement learning data. We report mean (and 95% HDIs) for the ‘shift’ parameters modeling cue-related effects, and Bayes Factors for directional effects (erotic>neutral). * (**): Strong evidence (dBF>10 /<.1) for a cue exposure effect (& 95% HDIs outside of zero).

|  | mean [95% HDI] | directional BF |
| --- | --- | --- |
| $s_{\eta_1}$ | .16 [-.78, 1.11] | 1.71 |
| $s_{\eta_2}$ | .29 [-.05, .65] | 19.15* |
| $s_{\gamma}$ | **-.31 [-.55, -.07]** | **.007\*\*** |
| $s_{\rho}$ | .07 [-.12, .29] | 2.87 |
| $s_{\beta_{mb}}$ | .33 [-6.08, 7.19] | 1.06 |
| $s_{\beta_{mf}}$ | -.01 [-1.89, 1.68] | .97 |
| $s_{\beta_2}$ | -.29 [-2.61, 2.18] | 0.69 |
| $s_{\alpha_1}$ | -.01 [-.07, .04] | .53 |
| $s_{\alpha_2}$ | 0.03 [-.01, .07] | 13.48* |
| $s_{\tau_1}$ | -.01 [-.03, .01] | .14 |
| $s_{\tau_2}$ | -.01 [-.03, .01] | .46 |
| $s_{v_{max1}}$ | **-.28 [-.47, -.1]** | **0.002\*\*** |
| $s_{v_{max2}}$ | .16 [-.13, .44] | 5.88 |

First, we confirmed the expected positive modulation of trial-wise drift rates by Q-value differences (i.e., the *β* weights for model-free, model-based, and S2 stage Q-values were reliably > 0, see figure 7b,c and *SI* - figure S4 e-g, all 95% HDIs > 0). In contrast to the regression analysis of participants’ S1 stay probabilities, but mirroring the S2 response time regression results, DDM showed no evidence for a reduction of model-based control (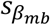, 95% HDI contained 0; figure 7b; table 3) following erotic vs. neutral cue exposure. Instead, we observed strong evidence for a reduction in the forgetting rate parameter (*s_γ_* = −.31, 95% HDI <0; figure 7a; table 3) modeling the forgetting rate of unchosen option values over time. However, in an exploratory correlation analysis, we found no significant association between changes in forgetting rate and the observed reduction in participants’ stay probabilities following small rewards and rare transitions (p=.46; figure 8a). In addition, the S1 drift-rate asymptote 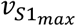, i.e. the maximum drift-rate modulation by value-differences, was reduced following erotic compared with neutral cue exposure (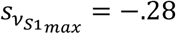, 95% HDI < 0; figure 7d; table 3). This may mirror an increased choice randomness as evident in participants’ overall reduced tendency of repeating the previous S1 choice following exposure to erotic cues. This was confirmed in an exploratory correlation analysis between the two measures (r=.79, p=8.07*10^−9, figure 8b). According to the 95% HDIs of the group-level posterior distributions, all other shift parameters showed no compelling evidence for a strong effect of erotic cue exposure. However, with respect to computed directional Bayes Factors (dBF), the learning rate for updating model-free Q-values in decision stage S2 (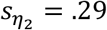, dBF=19.15; table 3, figure S3b) and the decision threshold in stage S2 (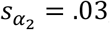, dBF=13.48; table 3, figure S3i) showed strong evidence for an increase following erotic vs. neutral cue exposure. This increase in decision thresholds in stage S2 corresponded with the observed increase in S2 response times following erotic vs neutral cues (r=.54, p=7.38*10^−4, figure 8c).

**Figure 7.**
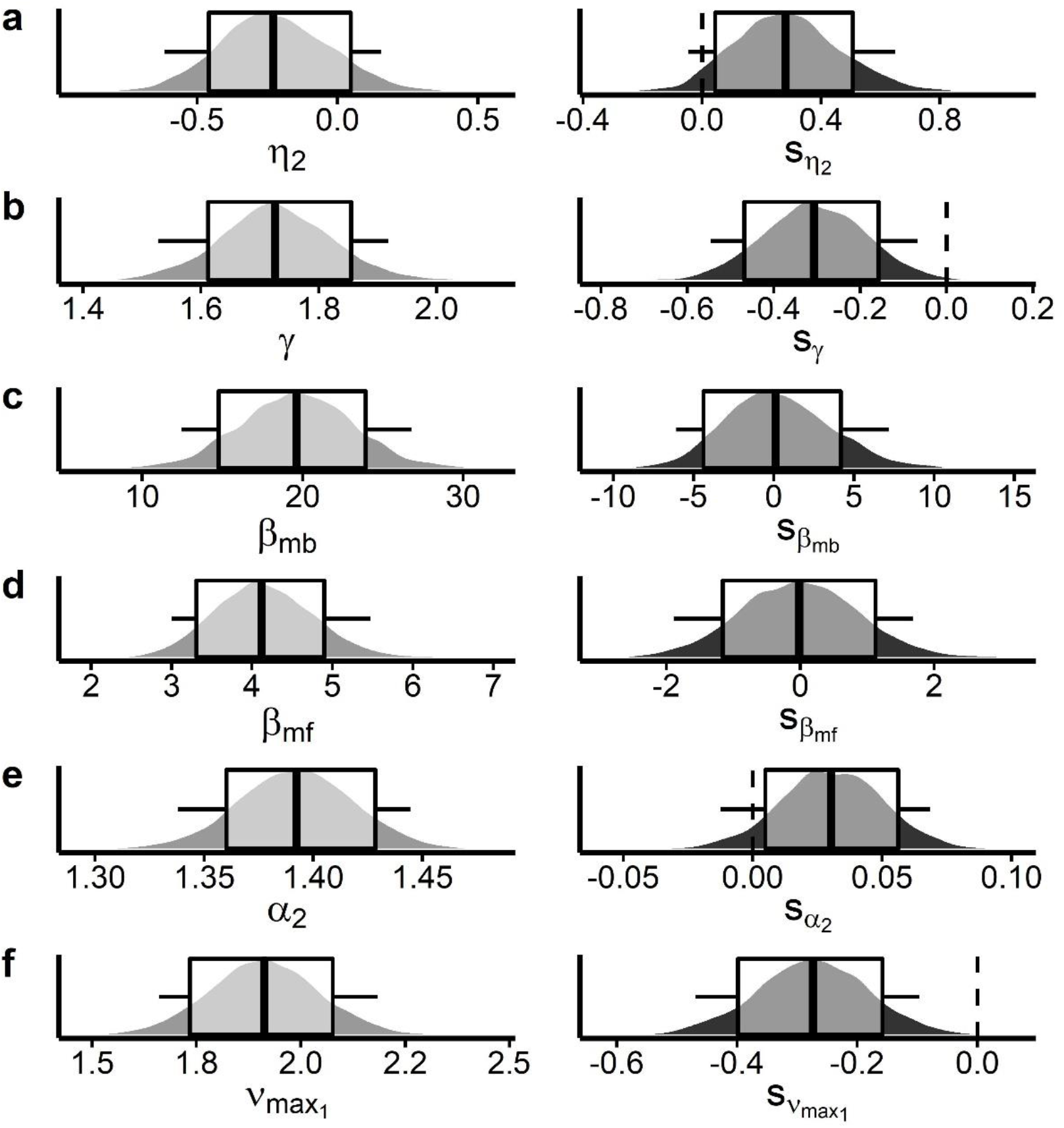
Selected posterior parameter distributions of the DDM_s_ group-level means of the sequential reinforcement learning data following neutral cue exposure (left column, grey plots) and their respective shifts as changes related to erotic vs. neutral cue exposure (right column, black plots). Boxplots depict 80% and 95% HDIs. Depicted parameters are (a) learning rate *η*_2_ for prediction error related updating in S2, (b) forgetting rate (forgetting rate) of unchosen option values (*γ*), (c) model-based parameter *β_mb_*, model-free parameter *β_mf_*, (e) decision threshold *α*_2_ and (f) S1 drift-rate asymptote 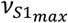. Only (b) the forgetting rate of unchosen options (*γ*) and (f) S1 drift-rate asymptote 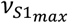 showed compelling evidence for negative shifts following erotic cue exposure according to their 95% HDIs (< 0). However, directional Bayes Factors showed evidence for additional positive shifts in (a) S2 learning rate *η*_2_ and decision threshold *α*_2_. We found no evidence for a shift in the parameters covering (b) model-based or (c) model-free reinforcement learning. (For inspection of all parameter distributions please see *SI*, figure S4. For inspection of parameter distributions from the softmax model formulation please see *SI* figure S5).

**Figure 8.**
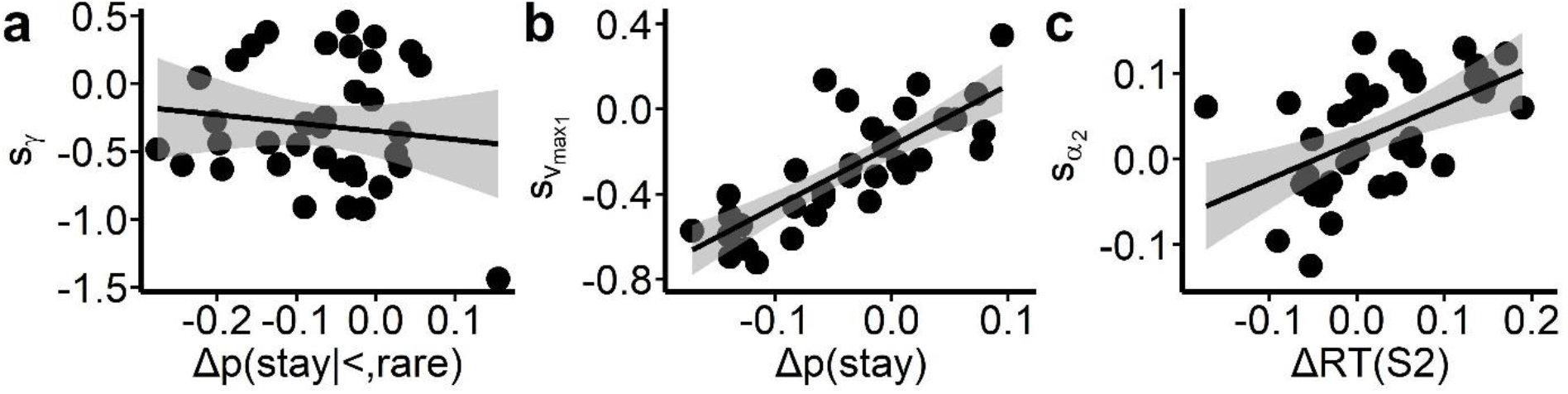
Exploratory correlation analysis results between observed model-agnostic findings of erotic cue exposure and findings from posterior distributions of the DDM_s_. (a) The shift in forgetting rate parameter (***s_γ_***) was not directly linked to changes in stay probabilities following smaller than expected rewards and rare transitions. (b) The shift in the maximum drift-rate modulation by value differences in stage S1 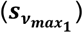 mirrored the general reduction in the tendency to repeat the last S1 choice (**Δ*p***(***stay***)). (c) The increase in response times in stage S2 was associated with a slight increase in decision thresholds 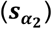.

### Correspondence between delay discounting and model-based control

It seems reasonable to assume that severity of impulsive choice behavior and strength of goal-directed control are related in a negative manner. However, when assessed via delay discounting and model-based control during reinforcement learning it seems that these measures are related to distinct cognitive processes (Solway et al., 2017). In an exploratory attempt, we tested if measures in both tasks are correlated. First, we tested whether proportion of LL choices and individual beta weights from the reward*transition interaction were correlated. There, was no significant association between both measures (r=0.19, p=.28). Likewise, we found no association of related model parameters, the individual discounting parameter *k* and model-based parameter *β_mb_* (r=−0.12, p=.48). A tendency for impulsive choices is also related to the starting point bias parameter *z*. We also did not observe a significant association between *z* and *β_mb_* (r=0.02, p=.92). In a similar manner, we tested whether changes in task performance measures following erotic cue exposure were associated. Individual reduction in LL choice proportions were not significantly correlated with individual beta weights from the reward*transition*condition interaction (r=0.09, p=0.63). However, the shift in the starting point bias *s_z_* during delay discounting was significantly correlated with the shift in the 2^nd^ stage decision threshold parameter 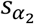 (r=−0.44, p=7.54*10^−3) during reinforcement learning, the remaining correlations with shifts in 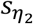, *s_γ_*, and 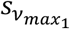 were all not significant (all p>=0.21).

## Discussion

Exposure to appetitive cues has long been discussed as an environmental driver of impulsive and maladaptive behavior (Kim & Zauberman, Wilson & Daly, usw.), but the underlying computational mechanisms as well as the generality of the effects across related cognitive constructs has remained elusive.

To address this, we investigated the effect of erotic cue exposure on two key aspects of value-based decision-making, temporal discounting and model-based reinforcement learning. In addition, we quantified self-reported and autonomic markers of arousal. We used drift diffusion modeling (DDM) in a hierarchical Bayesian approach to reliably quantify erotic cue exposure associated changes within distinct aspects of the dynamic choice processes in both tasks. Validating the overall approach, erotic cue exposure increased self-reported and autonomic arousal (pupil dilation, LF/HF ratio). Individual eye blink rate at baseline predicted arousal related changes in pupil dilation. In line with our predictions, erotic cues tended to increase discounting of delayed rewards. This was associated with a shift in the starting point bias towards smaller-sooner options in the DDM. For the first time, we report cue exposure effects on model-based reinforcement learning. Erotic cue exposure reduced signatures of model-based control in S1 stay probabilities. Interestingly, this effect was neither mirrored in cue exposure related changes of individual response time differences following rare vs. common transitions, nor in a change in model-based control parameter in the DDM analysis. Instead, cue exposure attenuated forgetting rates of unchosen option values. Taken together, across two prominent tasks, erotic cue exposure shifted behavior towards a less prospective decision mode driven by short-sighted strategies.

### Erotic cue exposure increases self-reported and autonomic arousal

To validate the experimental cue exposure manipulation, we assessed both self-reported and physiological arousal. Erotic cue exposure was associated with increased self-reported arousal as compared with neutral cue exposure. We quantified autonomic arousal via assessment of pupil dilation, heart rate and heart rate variability (low frequency to high frequency ratio, LF/HF) before cue exposure (baseline) and during erotic vs. neutral cue exposure. In addition, we assessed spontaneous eye blink rate as a potential proxy measure of central dopamine transmission (Jongkees and Colzato, 2016) at baseline.

We observed an increase in pupil dilation during exposure to erotic vs. neutral cues. Pupil dilation (Fotiou et al., 2000; Loewenfeld, 1999; Steinhauer et al., 2004) and heart rate (Berntson et al., 1997) are affected by both sympathetic and parasympathetic afferents. Recently, we reported increased phasic pupil dilation in response to erotic vs neutral cues in a trial-wise study design (Knauth and Peters, 2022). Here, we replicate this finding of increased pupil dilation during erotic vs. neutral cue exposure in a block-wise cue exposure design. Increased pupil dilation in response to arousing stimuli may stem from noradrenaline release from nucleus coeruleus (Abercrombie et al., 1988; Phillips et al., 2000; Murphy et al., 2011). As a result of a complex interplay of several nuclei within the brain (see e.g. Joshi and Gold, 2020), pupil dilation is controlled via sympathetic and parasympathetic inputs (Schumann et al., 2020). While pupil constriction is accomplished by parasympathetic innervation of the pupillae sphincter, pupil dilation is controlled by sympathetic innervation of the dilator pupillae muscle (Andreassi, 2000; Loewenfeld, 1999). Therefore, erotic cue exposure associated pupil dilation likely results from sympathetic effects. In an exploratory attempt, we found that individual eye blink rate at rest was predictive of the change in pupil dilation during erotic cue exposure. This corresponds with findings from Van Slooten et al. (2017) who reported that participants with lower eye blink rate showed stronger pupil dilation increases in response to reward expectation and surprise during reversal learning than participants with higher eye blink rate. In Parkinson’s disease patients, dopaminergic medication was shown to reinstate blunted pupil responses during reward expectation (Manohar et al., 2015). Taken together, this may hint at a potential dopaminergic role in the observed pupil dilation following erotic cue exposure and its association with eye blink rate.

In a recent event-related study Knauth and Peters (2022) found a nominal effect of cue exposure on heart rate 2-10 seconds following erotic cue exposure. We did not observe a modulation of heart rate during block-wise cue exposure. This likely relates to the block-wise design that hampers the detection of these so called ‘orienting responses’. However, we found evidence for an increased vagal (i.e. parasympathetic) relative to sympathetic input to cardiovascular activity during erotic vs. neutral cue exposure as indexed by attenuated LF/HF ratio of heart rate variability (Appelhans and Luecken, 2006). During exposure to emotional stimuli, prefrontal and anterior cingulate cortex might exert downstream modulation of parasympathetic cardiac innervation (Lane et al. 2009; Candia-Rivera et al., 2022) to adjust the cardiac pacemaker in relation to perceived motivational demands (Thayer and Lane 2009). In support of this account, we also found that higher self-reported arousal during erotic vs neutral cue exposure was associated with stronger attenuation of LF/HF ratio. Taken together, arousal effects verify that the present cue exposure manipulation yielded the predicted subjective and physiological effects on arousal. In particular our physiological results advance over previous studies that have focused solely on subjective measures (Wilson and Daly, 2004; Kim and Zauberman, 2013).

### Drift diffusion modeling

We implemented value-based decision-making (temporal discounting) and reinforcement learning models in a drift diffusion modeling (DDM) framework (Bruder et al., 2021; Fontanesi et al., 2019; Pedersen et al., 2017; Shahar et al., 2019; Wagner et al., 2020, 2022). Model comparison replicated previous results (Bruder et al., 2021; Fontanesi et al., 2019; Peters and D’Esposito, 2020; Wagner et al. 2020, 2022), such that data in both tasks were consistently better accounted for by models assuming a non-linear trial-wise drift rate modulation. Posterior predictive checks of choices and RTs confirmed the superior reproduction of relevant patterns (see *SI*). Prior parameter recovery work confirmed that DDM group-level parameters for temporal discounting data (Peters and D’Esposito, 2020; Wagner et al., 2020) and seq. reinforcement learning data (Shahar et al., 2019) recovered well. Notably, this modeling approach allowed us to link erotic cue exposure-related behavioral changes in both tasks to specific aspects of valuation and decision processes, namely a shift in the starting point bias (temporal discounting) and a shift in the forgetting rate of unchosen options (sequential reinforcement learning). This highlights the valuable insights that comprehensive modeling approaches can add beyond mere analysis of choices and RTs in neuro-cognitive research.

### Erotic cue exposure effects on temporal discounting

We observed a numerical enhancement of temporal discounting following erotic vs. neutral cue exposure as indexed by fewer larger-later choices. This concords with findings from previous studies that used block-wise exposure to appetitive (Li, 2008) and specifically erotic cues (Kim and Zauberman, 2013; Van den Bergh et al., 2008; Wilson and Daly, 2004) prior to task performance. Out-of-domain wanting for immediate reward in response to primary reinforcers such as erotic pictures has been ascribed to this effect (Van den Bergh et al., 2008). On the neural level, elevated tonic dopamine transmission in reward-related brain areas following sustained exposure to highly salient stimuli such as erotic pictures may be a potential underlying mechanism (O’Sullivan et al., 2011; Redouté et al., 2000). This view is supported by previous evidence showing that pharmacological modulation of central dopamine transmission impacts temporal discounting behavior (Petzold et al., 2019; Pine et al., 2010; Wagner et al., 2020). In addition to a potential role of dopamine transmission, arousal-related enhancement of noradrenaline release may likewise be associated with the modulatory effect of erotic cue exposure on discounting performance (Ventura et al., 2008). As reported in earlier studies (Finke et al., 2017; Kinner et al., 2017; Knauth and Peters., 2022) and as observed here, highly arousing cues increase pupil dilation, a measure associated with locus coeruleus activity and noradrenaline release (Aston-Jones and Cohen, 2005; Murphy et al., 2011), but see Megemont et al. (2022) for a more fine-graded analysis.

Diffusion modeling allowed us to disentangle effects of erotic cue exposure on the overall degree of temporal discounting (log(k)) and the starting point (bias) of the evidence accumulation process. Following erotic cue exposure, participant’s bias was substantially shifted towards the smaller-sooner option. This concords with the assumption of a bias towards out-of-domain wanting for immediate rewards (Van den Bergh et al., 2008) following exposure to erotic stimuli. Previous work in the perceptual decision-making domain has shown that increasing the value of one option in two alternative forced choice tasks shifts the starting point bias towards this option (Mulder et al., 2012; Fan et al., 2018). Optogenetic stimulation of striatal D1-receptor dopaminergic neurons in rodents during performance of a visual detection task enhanced approach behavior by positively shifting the expected value of visual events (Wang et al., 2018). In line, electrical stimulation of caudate nucleus neurons in monkeys resulted in a choice bias that was reflected in a shift of the starting point bias DDM parameter (Ding and Gold, 2012). Recently, Pedersen et al. (2021) linked reduced reward approach behavior in patients suffering from major depression to a shift in the starting point bias that was differentially modulated by nucleus accumbens (BOLD) activity compared with healthy controls. Taken together, this suggests that increased striatal dopamine signaling following erotic cue exposure as a candidate neural substrate of shifted preference for immediate reward options in temporal discounting.

### Erotic cue exposure effects on sequential reinforcement learning

In the model-agnostic analysis, we found that erotic vs. neutral cue exposure reduced participants’ tendency to stick with the last S1 choice in general. This may result from increased decision noise following erotic cue exposure. In the computational model relying on softmax action selection we observed that model-free and model-based coefficients were slightly reduced (see *SI*), an effect that reflects higher choice stochasticity. In a recent trial-wise cue exposure study, Knauth and Peters (2021) found that arousing cues seemed to increase participants’ decision noise to some extent although this effect was only apparent for aversive stimuli. In addition to a general cue effect on individual stay probabilities in S1, we observed that exposure to erotic vs. neutral cues resulted in attenuated model-based signatures in participants’ S1 stay probabilities. Post-hoc tests revealed that this was most evident following rare transitions and smaller-than-expected reward. Interestingly, individuals suffering from gambling disorder similarly showed a reduction in model-based control specifically after rare transitions and unrewarded trials (Wyckmans et al., 2019). They assumed that by attenuating inhibition within the dopamine D2-receptor related indirect pathway of the basal ganglia a putatively hyperdopaminergic state in pathological gamblers (Boileau et al., 2014; van Holst et al., 2018) may underly this specific reward expectancy violation associated model-based deficit. Although this is speculative, this resonates with the assumption that exposure to erotic vs neutral pictures may lead to heightened dopamine transmission within striatum causing the observed specific deficits in incorporating model-based information (transition structure of the task) following unexpected low reward outcomes. Notably, analysis of S2 response times following common vs. rare transitions showed no signature of attenuated model-based control. Here, we only observed slightly increased response times in general. This clearly speaks against a plain enhancement of motor impulsivity following erotic vs. neutral cue exposure.

Unexpectedly, diffusion modeling did not link the attenuated signatures of model-based control in S1 stay probabilities to a reduction in a model-based drift-rate modulation 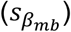. Instead, erotic cue exposure affected more basic processes related to evidence accumulation in S1 (maximum drift rate) and value tracking (forgetting rates). Notably, following initial evidence for a direct impact of dopamine on the balance between model-based and model-free control (Wunderlich et al., 2012; Sharp et al., 2016), recent work challenged this view (Kroemer et al., 2019). We observed reduced drift-rate asymptotes in S1, which were correlated strongly with a general reduction in participants’ tendency to repeat the last choice in S1. This observation is in line with the idea that exposure to erotic cues increases decision noise, as has been shown previously for aversive stimuli (Knauth and Peters, 2022). Such modulations of choice stochasticity might likewise be related to increased striatal dopamine signaling (Zhuang et al., 2001; Kayser et al., 2015; Chakroun et al., 2020).

This effect was accompanied by strong evidence for attenuated forgetting rates (*s_γ_*) i.e. increased forgetting of unchosen Q-values following erotic cue exposure. Forgetting of expected values in reinforcement learning tasks has been linked to specific aspects of striatal dopamine signaling, so called ramping (Morita and Kato, 2015). Thus, increased tonic dopamine levels within striatum following erotic cue exposure may have interfered with this phenomenon e.g. via enhanced D2-receptor activation, although this is highly speculative.

Directional Bayes factors showed stronger evidence for an increase vs. a decrease in the updating of model-free Q-values in S2 (positive shift in the S2 learning rate following erotic cue exposure, 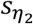). Model-free prediction error-related updating of Q-values is tightly linked to dopamine (e.g. Schultz et al., 1997; Daw et al., 2011) which also speaks for a potential role of increased dopamine signaling following erotic cue exposure as an at least partly underlying neural mechanism of the observed alterations in sequential reinforcement learning performance. Finally, in accordance with the observed slowing of S2 response times in the model-agnostic analysis, we found stronger evidence for an increase rather than a decrease in decision-thresholds in S2. The decision threshold parameter within a DDM framework has been linked to subthalamic nucleus activity (Aron et al., 2016; Herz et al., 2016) that is under dopaminergic and noradrenergic control (Canteras et al.,1990; Cragg et al., 2004; Parent and Hazrati, 1995).

### Correspondance between delay discounting and model-based control

Similar to a previous study in an online cohort (Solway et al., 2017), we found no evidence for an association of delay discounting and model-based control during reinforcement learning. This seems at odds with the observation that both measures show alterations in people suffering from addiction (Bickel et al., 2019; Lucantonio et al., 2014), and that modulation of dopamine transmission (e.g. via L-Dopa or Haloperidol) intake modulates both model-based control (Wunderlich et al., 2010; Sharp et al., 2016) and temporal discounting (de Wit et al., 2002; Pine et al., 2010; Wagner et al., 2020). Recently, we found that a single dose of the catecholamine precursor L-Tyrosine resulted in faster response times in both tasks due to reduced decision thresholds according to DDM (Mathar et al., 2022). However, the findings regarding dopaminergic manipulation of both model-based control (Kroemer et al., 2019; Deserno et al., 2021) and delay discounting (Petzold et al., 2019) are still heterogeneous. Interestingly though, the shift in the starting point bias towards impatient choices in the delay discounting task following erotic cue exposure was associated with an increase in the speed-accuracy trade-off (shift in the decision-threshold parameter) in the 2^nd^ decision stage of the seq. reinforcement learning task. The 2^nd^ decision stage is related to pure model-free prediction error updating of respective choice options. Possibly, erotic cue exposure rendered these immediate reward values more salient similar to the smaller sooner choice options in the delay discounting task which resulted in more cautious evidence accumulation in this model-free valuation related decision stage.

### Limitations

Our study has a few limitations that need to be acknowledged. First, we only included male participants. To date, erotic cue exposure effects on reward-based decision making such as temporal discounting have been primarily revealed in male volunteers (Kim and Zauberman, 2013; Van den Bergh et al., 2008; Wilson and Daly, 2004). Women and men may differ in their physiological response to arousing stimulus material (Bradley et al., 2001; Lithari et al., 2010; Wrase et al., 2003; Finke et al., 2017), although a recent meta-analysis found no evidence for different neural substrates of sexual arousal (Mitricheva et al., 2019). Future studies should therefore extend the present approach and include participants from both sexes. Second, although self-reported arousal differences between erotic and neutral cue exposure appeared to be strong, the differences in autonomic arousal were small in comparison and warrant a replication in a larger sample size. Third, while high test-retest reliabilities of temporal discounting measures are well established in the literature (Kirby 2009; Peters and Büchel, 2009; Bruder et al., 2021 usw.), the reliability of model-based behavior on the 2-step task has recently been called into question (Enkavi et al., 2019). Fortunately, despite the exposure manipulation, we found good test-retest reliability of participants’ individual model-based control during reinforcement learning between sessions that was considerably higher than reported in Enkavi. This may stem from slight task design changes in relation to the standard task version (Daw et al., 2011). We modified the outcome stage by replacing the fluctuating reward probabilities (reward / no reward) with fluctuating reward magnitudes (between 0 and 100) (see Kool et al., 2016). Likewise, participants completed 300 trials instead of the more common 201 trials.

## Conclusion

Here, we show that exposure to erotic stimuli increased self-reported arousal and modulated autonomic nervous system activity as indexed by pupil dilation and heart rate variability. Strikingly, erotic cue exposure modulated task performance in two distinct but conceptually related reward-based decision-making tasks considered to index trans-diagnostic constructs. First, we replicate previous work of increased temporal discounting and second, for the first time we report alterations in model-based reinforcement learning following erotic cue exposure. Hierarchical Bayesian drift diffusion modeling (DDM) revealed that enhanced discounting resulted from a shift in the starting point bias towards impatient choice options. Reduced model-based signature in participants’ stay-shift choice patterns was not mirrored in an attenuated DDM parameter covering model-based reinforcement learning strategy but resulted from differential changes in forgetting rates of unchosen option values and reduced drift-rate asymptotes. Thus, exposure to erotic cues seemed to shift decision-making away from a more prospective, goal-directed mode towards choices related to more immediate rewards via specific alterations in valuation and model-free reinforcement learning related processes. Our work highlights the deeper insights psychophysiological research can gather from sophisticated computational modeling approaches.

## Acknowledgements

This work was funded by Deutsche Forschungsgemeinschaft (PE1627/5-1 to J.P.)

## Supplementary Information (SI)

### Demographic & psychological screening

**Table S1.** Study sample characteristics (N=39). SE= standard error; BMI: body mass index; YOE=years of education; (monthly) income; average amount of sleep (hours/night of the past 4 weeks), BDI-II: Beck Depression Inventory-II; BIS-15: Barratt-Impulsiveness Scale (15 items); BIS/BAS: Behavioral Inhibition and Behavioral Activation System.

|  | mean (range) | SE |
| --- | --- | --- |
| age | 25.23 (19-34) | 0.65 |
| BMI | 24.00 (19.44-33.14) | 0.40 |
| YOE | 12.78 (8-30) | 0.49 |
| income | 1099.33 (0-3500) | 137.86 |
| sleep quantity | 7.23 (5-9) | 0.13 |
| BDI-II | 6.38 (0-17) | 0.78 |
| BIS-15 | 31.56 (21-45) | 1.04 |
| BIS(BIS/BAS) | 2.60 (1.57-3.29) | 0.44 |
| BAS(BIS/BAS) | 3.13 (2.08-3.69) | 0.66 |

### Model-comparison, predictive accuracy

Depicted below is the model fit comparison according to Watanabe-Akaike Information Criterion (WAIC) and the estimated log pointwise predictive density (elpd).

**Table S2.** Model fit comparison of the DDMs to a DDM with a linear modulation (DDM_lin_) and without modulation (DDM_0_) of drift rate by value differences in the temporal discounting and the seq. reinforcement learning task via the Watanabe-Akaike Information Criterion (WAIC), the estimated log pointwise predictive density (elpd), and its difference to the winning model (DDM_s_).

|  | WAIC | -elpd | -Δelpd | 95% CI (-Δelpd) |
| --- | --- | --- | --- | --- |
| Temp. Discount. |  |  |  |  |
| **DDM<sub>0</sub>** | 20816 | 10466 | 3080 | 2778 - 3381 |
| **DDM<sub>lin</sub>** | 16245 | 8185 | 799 | 624 - 974 |
| **DDM<sub>s</sub>** | 15253 | 7386 | - | - |
| Seq. RL |  |  |  |  |
| **DDM<sub>0</sub>** | 29198 | 14664 | 10905 | 9743 - 12068 |
| **DDM<sub>lin</sub>** | 8644 | 4387 | 628 | 463 - 794 |
| **DDM<sub>s</sub>** | 7405 | 3759 | - | - |

We compared the respective model formulations also in terms of predictive accuracy of participants’ binary choices in each task. While the softmax model formulation performed best, as it is only fitted to participants’ choices (and not to RT distributions), the DDM_s_ performed on a comparable level and better than the DDM_0_ and DDM_lin_ (Table S3).

**Table S3.** Proportions of correctly predicted binary choices (mean (range)) for the temporal discounting (TD) task data and for both stages (S1, S2) of the seq. RL choice data, respectively.

|  | softmax | DDM<sub>0</sub> | DDM<sub>lin</sub> | DDM<sub>s</sub> |
| --- | --- | --- | --- | --- |
| **TD task** | 90 (79-95) | 59 (50-95) | 75 (70-88) | 84 (76-95) |
| **seq. RL, S1** | 76 (49-94) | 54 (50-75) | 69 (50-87) | 72 (50-90) |
| **seq. RL, S2** | 82 (60-93) | 51 (50-52) | 73 (53-85) | 77 (53-88) |

Values are computed via simulations based on 500 samples drawn from each of the respective single subject parameters’ posterior distributions.

### Posterior predictive checks

Posterior predictive checks (see ’Methods’ section for details) showed that the DDM_s_ reproduced the effect of decision conflict (value differences of choice options) on participants’ RTs and choice patterns best in comparison to alternative model formulations for both the temporal discounting data (figure S1) and for the seq. RL task (figure S2).

### Temporal discounting task

**Figure S1.**
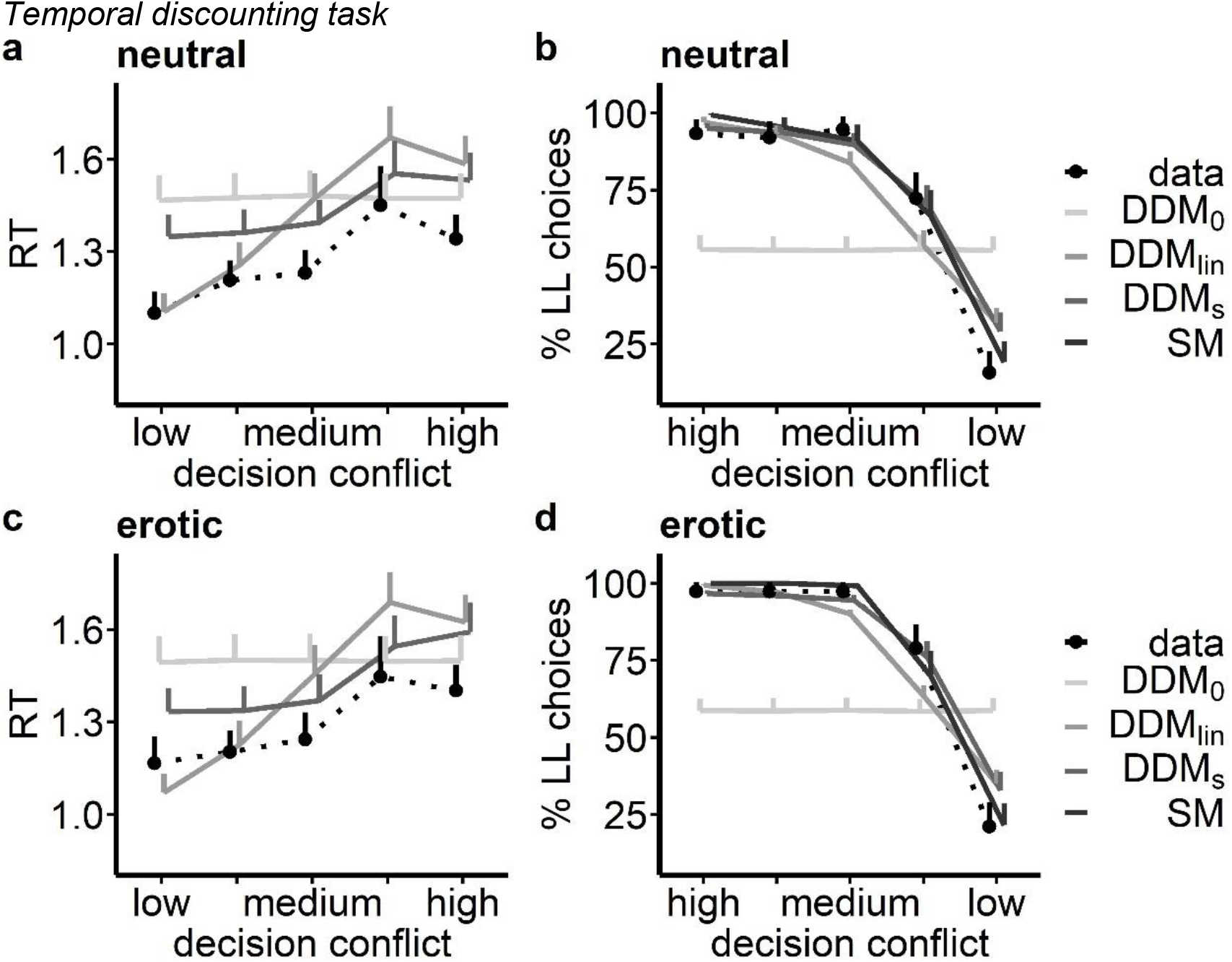
Posterior predictive checks for the winning DDM_s_ and alternative model formulations for the temporal discounting task data following neutral and erotic cue exposure. In each plot the dotted line depicts participants’ median RTs or mean % LL choices. Solid lines depict the median RTs or mean LL choices drawn from 500 simulations of each of the different DDM formulations, as well as of a standard softmax model for choice data. Data and simulations are plotted in relation to the absolute difference of LL (subjective values) and SS choice options.

### Seq. reinforcement learning task

**Figure S2.**
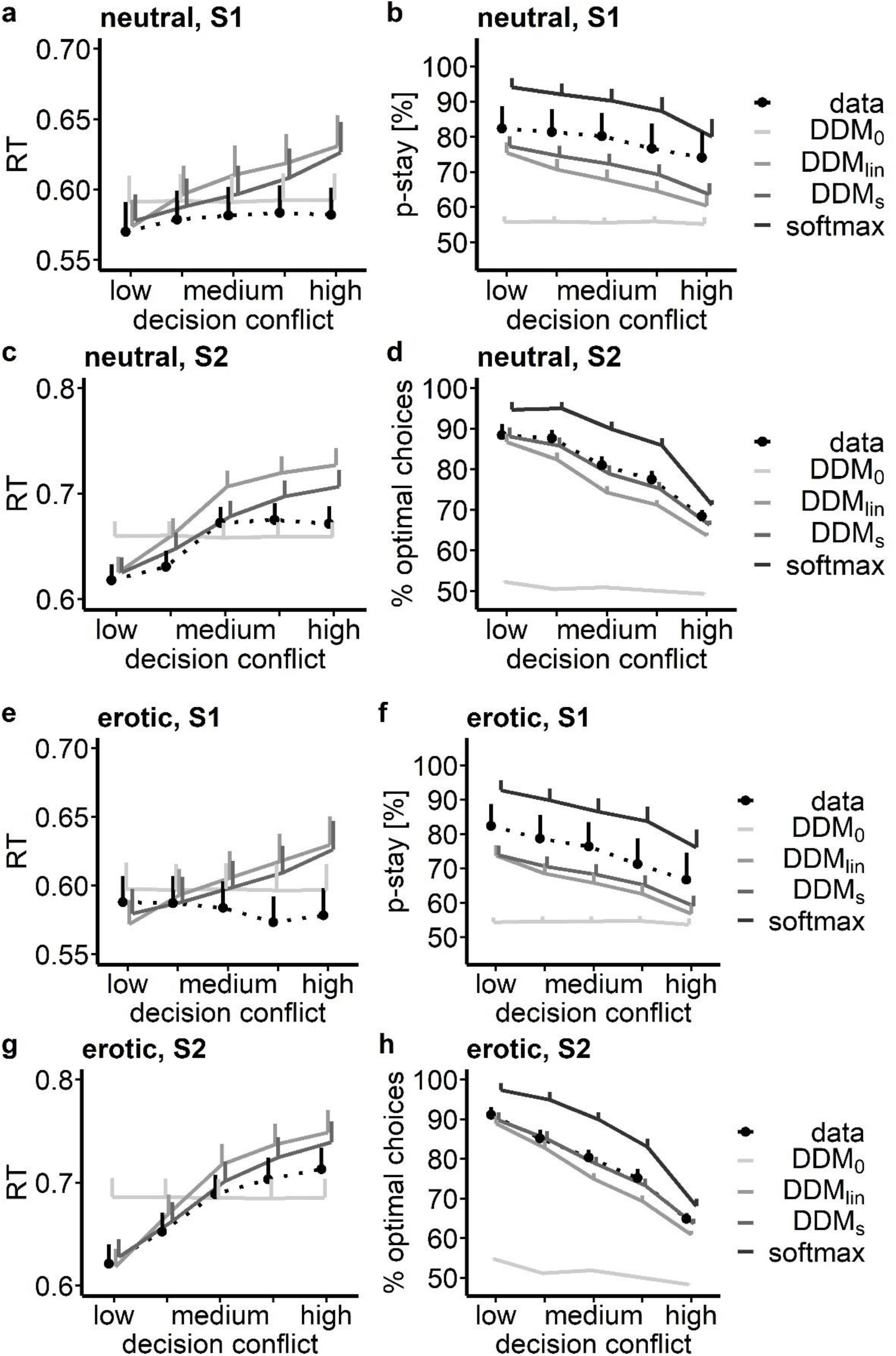
Posterior predictive checks for the winning DDM_s_ and alternative model formulations for the seq. RL task data following (a) – (d) neutral and (e)-(h) erotic cue exposure. In each plot the dotted line depicts participants’ median RTs or mean choice behavior. Solid lines depict the median RTs and mean choice behavior drawn from 500 simulations of each of the different DDM formulations, as well as of a standard softmax model for choice data. The upper row depicts RT data and simulations and participants’ probability to choose the same action as in the previous trial for the first decision stage S1 in relation to the options value differences (‘decision conflict’). The lower row depicts S2 RT data and fraction of optimal choices in S2 (highest value option chosen) of participants and related model simulations.

### Temporal discounting task, softmax model

We also modeled choice data in the temporal discounting task using a hyperbolic discounting model with standard softmax-action selection (see methods section). In this model, we found a nominal increase in discount rate log(k) following erotic cue exposure (mean [95% HDI]: *s_k_*=0.15 [-.09, 0.38], BF=8.36; figure S3). We found no evidence for a modulatory effect of cue exposure on modeled choice stochasticity (mean [95% HDI]: *s_β_*=0.02 [−0.03, 0.08], figure S3).

**Figure S3.**
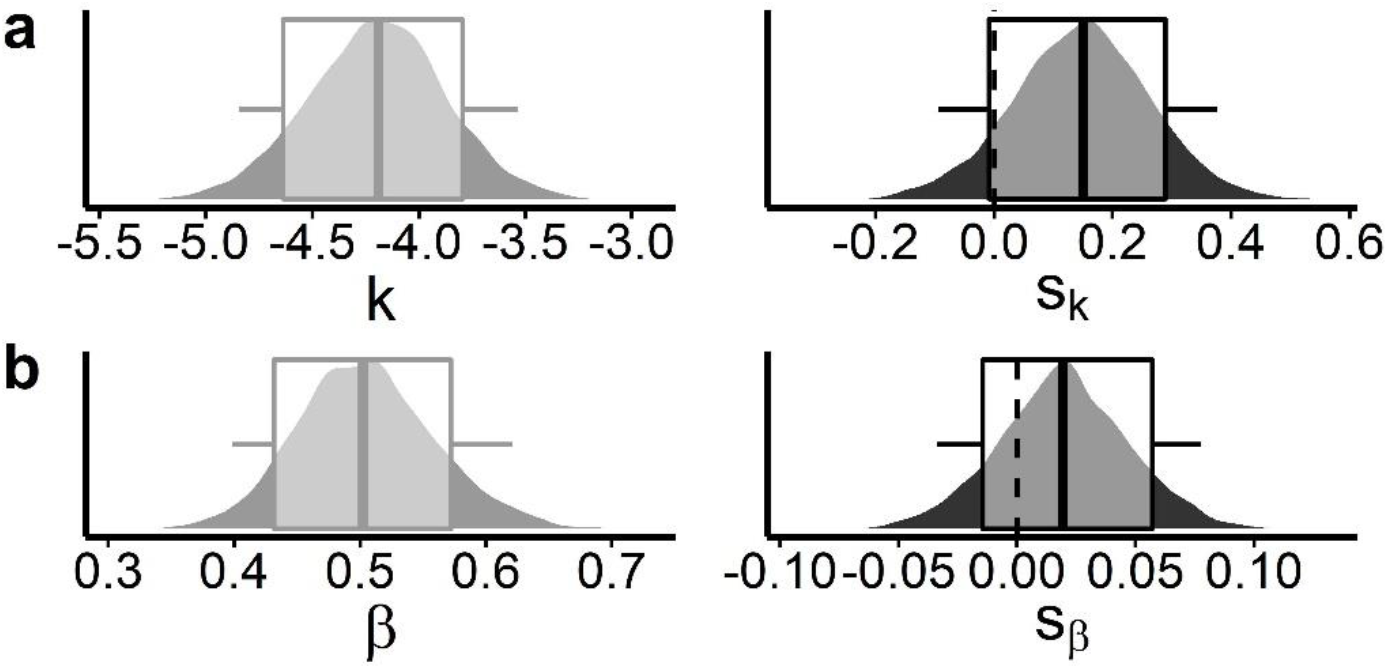
Posterior distributions of the group-level means of the softmax model of the temporal discounting task data following neutral cue exposure (a,b, grey plots) and erotic cue exposure related changes in these (c,d, black plots). Boxplots depict 80% and 95% HDIs. Depicted parameters are: discount rate log (*k*), softmax inverse temperature *β*, and erotic cue exposure related shifts thereof *s_k_*, *s_β_*.

### Sequential RL task, drift diffusion model

Figure S4 depicts the posterior distributions of all group-level mean parameters from the DDM_s_ following neutral cue exposure and their related shifts following erotic cue exposure.

**Figure S4.**
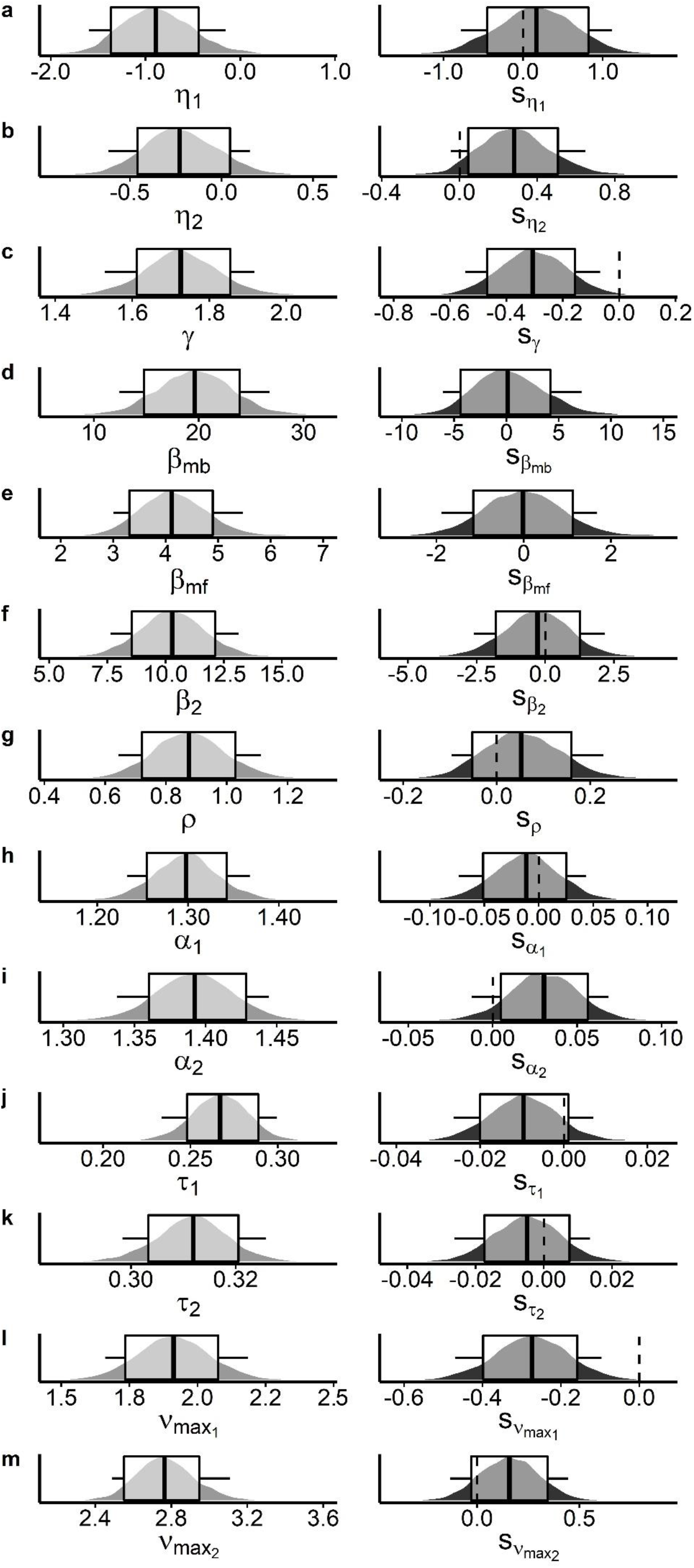
Group-level mean posterior distributions of all DDM_s_ model parameters following neutral cue exposure (left column, grey plots) and related shifts of these following erotic cue exposure (right column, black plots) for the seq. RL task data. Boxplots depict 80% and 95% highest density intervals (HDIs). Depicted parameters (left) and related shifts (right) are (a)-(c) learning-rates in decision stages S1 and S2 *η*_1_, *η*_2_, and decay rate of unchosen options *γ*; (d)-(f) weights for model-based Q-values *β_mb_*, weights for model-free Q-values *β_mf_* and weights for S2 stage Q-values *β*_2_; (g) choice perseveration parameter *ρ*; (h), (i) decision-thresholds for both stages *α*_1_, *α*_2_; (j), (k) non-decision times *τ*_1_, *τ*_2_ and (l), (m) the asymptotes of respective drift-rates 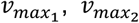.

### Sequential RL task, softmax model

We also implemented a similar computational reinforcement learning model that is based on softmax action selection instead of a DDM implementation (see Methods section). Note that our model formulation includes the nested versions with only one learning rate, no decay rate, no perseveration, no model-based and or no model-free effects as special cases, where the respective posteriors are centered at 0. Kruschke (2011) suggests to examine the posterior distributions of the full model in such cases, rather than performing a model comparison across all nested versions. The softmax model yielded comparable results to the DDM_s_. Erotic cue exposure resulted in strong evidence for a reduction in the forgetting rate parameter (*s_γ_*= −0.33, 95% HDI <0; figure S5c; table S4) modeling the forgetting rate of unchosen option values over time. According to the 95% HDIs of the group-level posterior distributions of the softmax model, all other shift parameters showed no compelling evidence for a strong effect of erotic cue exposure. However, with respect to computed directional Bayes Factors (dBF), and similar to the posterior distributions from the DDM_s_, the learning rate for updating model-free Q-values in decision stage S2 seemed increased following erotic cue exposure (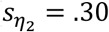, dBF=19.15; table S4, figure S5b). In contrast to the DDM_s_ findings, there was also moderate evidence for a reduction in model-based control according to dBF (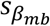 = −0.43, dBF=0.10, table S4).

**Table S4.** Erotic cue exposure associated changes in group-level means of the softmax model for the sequential reinforcement learning data. We report mean (and 95% HDIs) for the ‘shift’ parameters modeling cue-related effects, and Bayes Factors for directional effects (erotic>neutral). * (**): Strong evidence according to dBF>10 /<.1 (& 95% HDIs outside of zero) for a cue exposure effect.

|  | mean [95% HDI] | directional<br>BF |
| --- | --- | --- |
| $s_{\eta_1}$ | 0.67 [-0.34, 1.57] | 1.71 |
| $s_{\eta_2}$ | 0.30 [-0.04, 0.67] | 19.15* |
| $s_\gamma$ | **-0.33 [-0.49, -0.18]** | **2.19\*10<sup>-6</sup>\*\*** |
| $s_\rho$ | -0.05 [-0.17, 0.06] | 0.24 |
| $s_{\beta_{mb}}$ | -1.51 [-3.64, 0.64] | 0.10* |
| $s_{\beta_{mf}}$ | -0.43 [-1.16, 0.40] | 0.16 |
| $s_{\beta_2}$ | 0.87 [-0.58, 2.53] | 7.27 |

**Figure S5.**
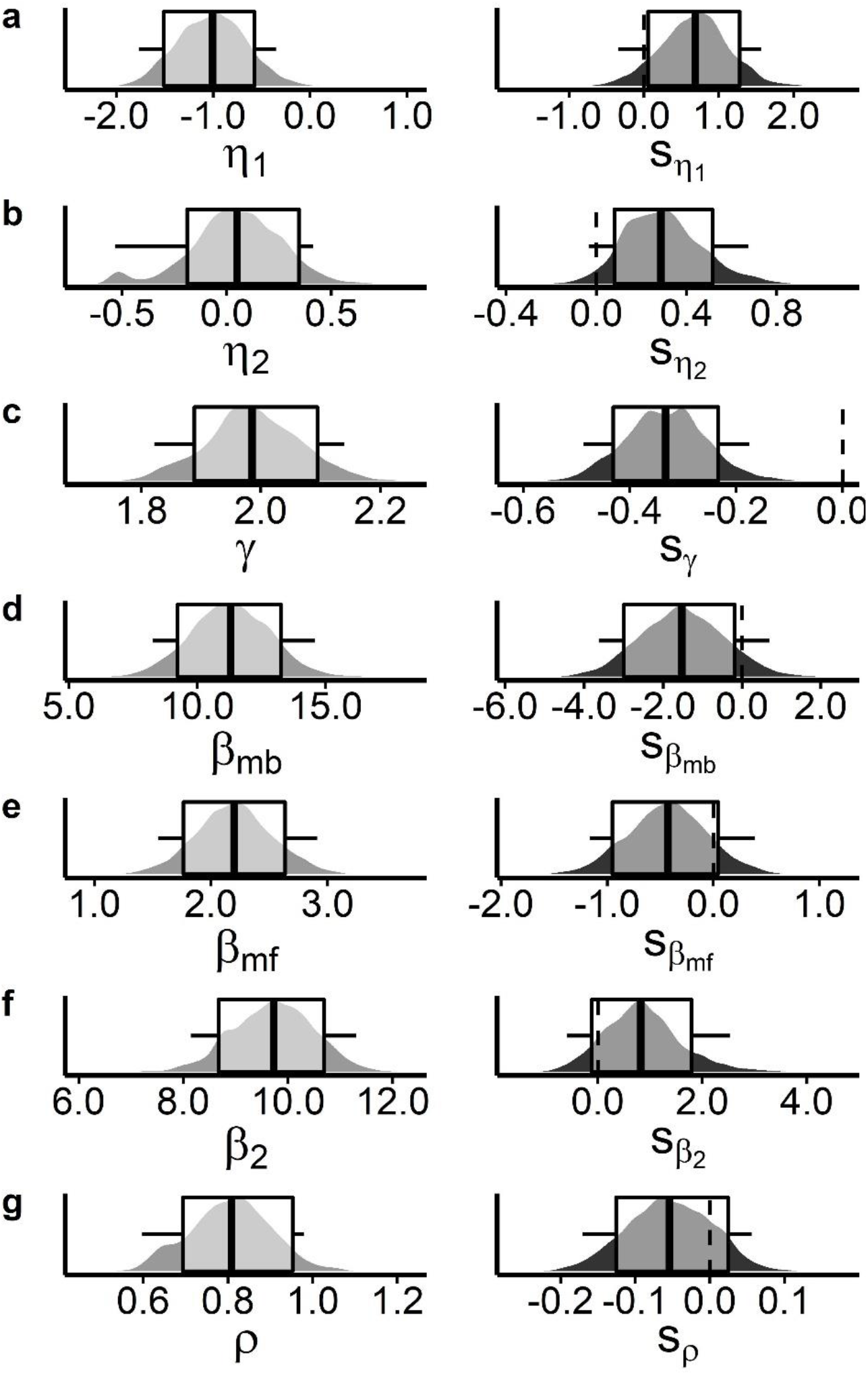
All group-level mean posterior distributions of the softmax model parameters following neutral cue exposure (left column, grey plots) and related shifts of these following erotic cue exposure (right column, black plots) for the seq. RL task data. Boxplots depict 80% and 95% highest density intervals (HDIs). Depicted parameters (left) and related shifts (right) are (a)-(c) learning-rates in decision stages S1 and S2 *η*_1_, *η*_2_, and decay rate of unchosen options *γ*; (d)-(f) weights for model-based Q-values *β_mb_*, weights for model-free Q-values *β_mf_* and weights for S2 stage Q-values *β*_2_ and (g) choice perseveration parameter *ρ*.

